# Transcription factors orchestrate dynamic interplay between genome topology and gene regulation during cell reprogramming

**DOI:** 10.1101/132456

**Authors:** Ralph Stadhouders, Enrique Vidal, François Serra, Bruno Di Stefano, François Le Dily, Javier Quilez, Antonio Gomez, Samuel Collombet, Clara Berenguer, Yasmina Cuartero, Jochen Hecht, Guillaume Filion, Miguel Beato, Marc A. Marti-Renom, Thomas Graf

## Abstract

Chromosomal architecture is known to influence gene expression, yet its role in controlling cell fate remains poorly understood. Reprogramming of somatic cells into pluripotent stem cells by the transcription factors (TFs) Oct4, Sox2, Klf4 and Myc offers an opportunity to address this question but is severely limited by the low proportion of responding cells. We recently developed a highly efficient reprogramming protocol that synchronously converts somatic into pluripotent stem cells. Here, we employ this system to integrate time-resolved changes in genome topology with gene expression, TF binding and chromatin state dynamics. This revealed that TFs drive topological genome reorganization at multiple architectural levels, which often precedes changes in gene expression. Removal of locus-specific topological barriers can explain why pluripotency genes are activated sequentially, instead of simultaneously, during reprogramming. Taken together, our study implicates genome topology as an instructive force for implementing transcriptional programs and cell fate in mammals.

## INTRODUCTION

Somatic cell reprogramming into pluripotent stem cells (PSCs) represents a widely studied model for dissecting how transcription factors (TFs) regulate gene expression programs to shape cell identity^1,2^. Chromosomal architecture was recently shown to be cell type-specific and critical for transcriptional regulation^3-5^, but its importance for cell fate decisions remains poorly understood.

Two major levels of topological organization have been identified in the genome^6-8^. The first level segregates the genome, at the megabase scale, into two subnuclear compartments. The A compartment corresponds to active chromatin typically associated with a more central nuclear position, while the B compartment represents inactive chromatin enriched at the nuclear periphery/lamina^9-14^. Compartmentalization is consistent amongst individual cells and a potential driver of genome folding^15^. A second sub-megabase level consists of topologically associated domains (TADs)^16-18^ and chromatin loops^11^, which restrict or facilitate interactions between gene regulatory elements^19,20^. Importantly, modifying chromatin architecture can lead to gene expression changes^19,21-24^. Moreover, *de novo* establishment of TAD structure during zygotic genome activation in Drosophila embryos has been shown to be independent of ongoing transcription, demonstrating that chromatin architecture is not simply a consequence of transcription^25^. Genome topology could therefore be instructive for gene regulation^26,27^, but whether this reflects a general principle that occurs on a genome-wide scale in space and time is unknown.

Mechanistic studies with mammalian cell reprogramming systems have been hampered by the typically small percentage of responding cells^1,28^. To overcome this shortcoming, we recently developed a highly efficient and synchronous reprogramming system based on the transient expression of the TF C/EBPα prior to induction of the Yamanaka TFs Oct4, Sox2, Klf4 and Myc (OSKM)^29,30^. OSKM activates the endogenous core pluripotency TFs sequentially in the order of Oct4, Nanog and Sox2, implying that locus-specific barriers dictate gene activation kinetics^31-33^. Here, we studied how C/EBPα and OSKM affect genome topology, the epigenome and gene expression during reprogramming. We found that the TFs bind hotspots of topological reorganization at both the compartment and TAD levels, exploiting existing 3D genome landscapes. Dynamic reorganization of genome topology frequently preceded gene expression changes at all levels and provided an explanation for the sequential activation of core pluripotency genes during reprogramming. Together, our observations indicate that genome topology has an instructive role in implementing transcriptional programs relevant for cell fate decisions in mammals.

## RESULTS

### TFs prime the epigenome for reprogramming

We exposed bone marrow-derived pre-B cells to the myeloid TF C/EBPα to generate ‘Bα cells’, which resemble granulocyte/macrophage progenitors^30^. The subsequent activation of OSKM induces the reprogramming of nearly 100% of Bα cells into PSC-like cells within 4-8 days^29,30^. To obtain a high-resolution map of changes in gene expression and chromatin structure we examined 6 different cell stages (B, Bα, D2, D4, D6, and D8) during reprogramming, as well as PSCs (**Fig.1a**). We profiled the transcriptome by RNA-Seq, active chromatin deposition by H3K4Me2 ChIPm-Seq^34^, and chromatin accessibility by ATAC-Seq^35^ (**Supplementary Fig.1**). Expression of half of all genes was significantly affected (FDR<0.01) between any two time points, starting with the rapid silencing of the core B cell program initiated by C/EBPα. Pluripotency genes were then activated sequentially, with the core pluripotency factors Oct4, Nanog and Sox2 being activated at D2, D4 and D6, respectively (**Fig.1b-c**).

**Figure 1.**
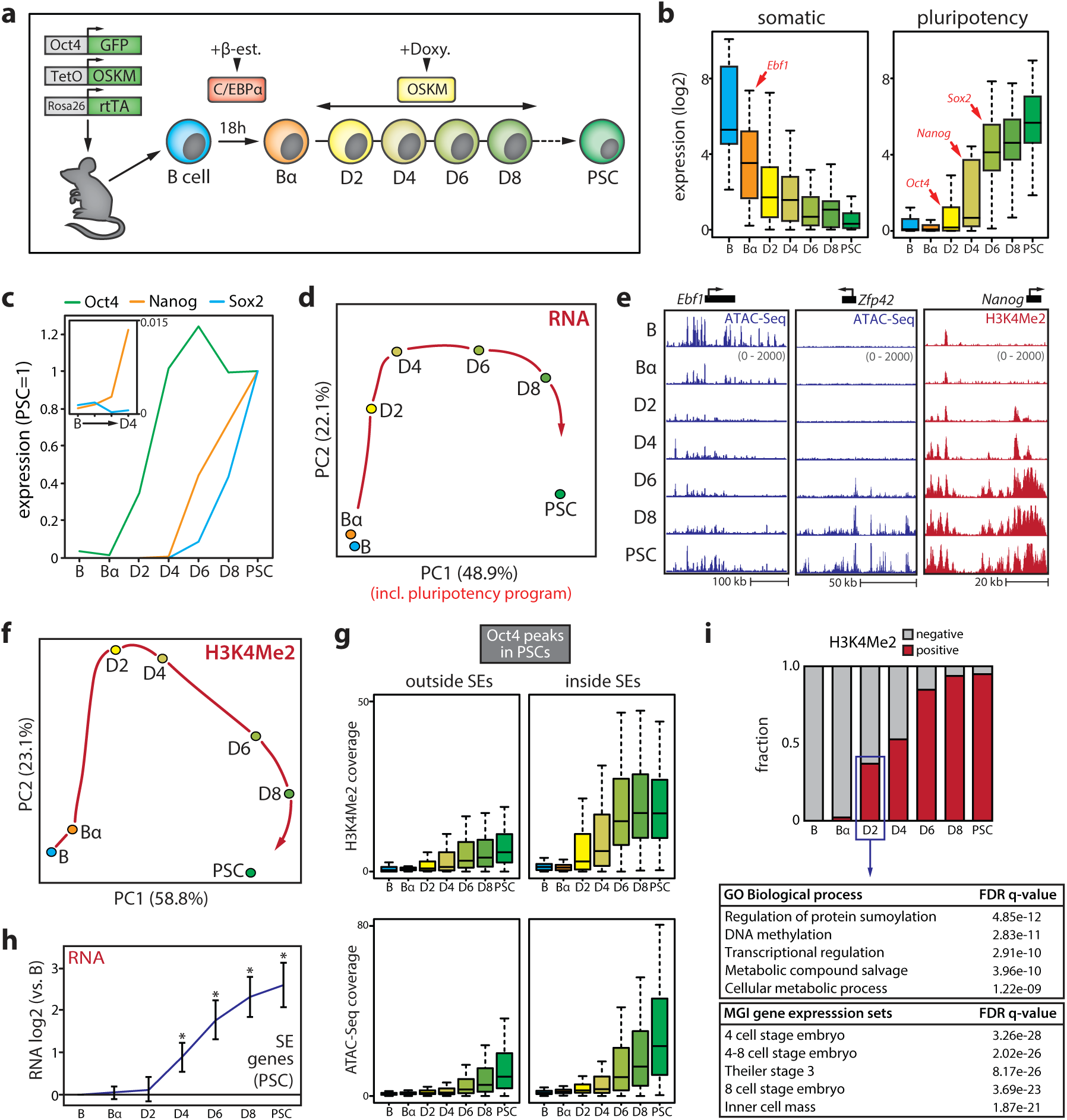
Dynamics of the transcriptome and epigenome during reprogramming. (**a**) Schematic overview of the reprogramming system. C/EBPα-ER in B cells is translocated into the nucleus upon beta-estradiol (β-est.) treatment. After β-est. wash-out, Oct4, Sox2, Klf4 and Myc (OSKM) are induced by doxycycline (doxy.). (**b**) Box plots of gene expression dynamics (normalized counts) of a set of core B cell (‘somatic’, n=25) and PSC (‘pluripotency’, n=25) identity genes. (**c**) Gene expression kinetics of Oct4, Nanog and Sox2 during reprogramming (relative to the levels in PSCs). Inset shows Nanog expression first appears at D4. (**d**) Principal component analysis (PCA) of gene expression dynamics during reprogramming. A red arrow indicates hypothetical trajectory. (**e**) Examples of chromatin opening (measured by ATAC-Seq) and H3K4Me2 deposition (measured by ChIPm-Seq) at gene regulatory elements controlling B cell (Ebf1) or pluripotency (Zfp42 and Nanog) genes. (**f**) PCA of H3K4Me2 dynamics during reprogramming. A red arrow indicates hypothetical trajectory. (**g**) Box plots of dynamics of H3K4Me2 deposition (top) and chromatin accessibility (bottom) at Oct4 binding sites outside and inside PSC superenhancers (SEs). (**h**) Expression dynamics of genes associated with a super enhancer (SE) in PSCs (error bars denote 95% CI, *p<0.01 expression versus B cells, unpaired two-tailed *t*-test). (**i**) Fraction of H3K4Me2^+^ Oct4 binding sites in PSC SEs during reprogramming; table shows a gene ontology (GO) analysis for genes associated with early Oct4 recruitment.

Principal component analysis (PCA) revealed a trajectory along which B cells acquire a PSC gene expression program (**Fig.1d**). Epigenome remodeling showed similar dynamics, with an early loss of chromatin accessibility at gene regulatory elements controlling the B cell program induced by C/EBPα followed by the establishment of active and open chromatin at pluripotency genes by OSKM (**Fig.1e, Supplementary Fig.1**). OSKM induction led to a genome-wide expansion of active chromatin marked by H3K4Me2, known to be deposited at both primed and active gene regulatory elements^36^ (**Supplementary Fig.1f**). The H3K4Me2 landscape more rapidly converged on a pluripotent state than gene expression, suggesting that OSKM primes regulatory elements for subsequent gene activation (**Fig.1f**). Many regions bound by Oct4 in PSCs^37^ had already acquired H3K4Me2 by D2, especially those located in superenhancer (SE) elements^37^ (**Fig.1g, Supplementary Fig.1i**). Chromatin opening occurred progressively at Oct4 binding sites (**Fig.1g, Supplementary Fig.1g-h**). 37% of Oct4 binding sites in PSC-SEs had already acquired an active chromatin signature by D2 (**Fig.1i**), while activation of most associated genes occurred 2 days later (**Fig.1h**). These early targeted SEs are linked to genes involved in DNA methylation (e.g. Tet1/2, Idh2), (post)-transcriptional regulation (e.g. Oct4, Nanog, Klf9) and metabolism (e.g. Upp1, Uck2), a gene signature strongly associated with 4 to 8 cell stage embryos (**Fig.1i**).

### Chromatin state, topology and transcription are dynamically coupled

We used in-situ Hi-C^11^ to map 3D genome organization during cell reprogramming at high resolution (**Supplementary Fig.2a**, see Supplemental Materials) and determined genome segmentation into A and B compartments (**Fig.2a**). Quantitative changes in A/B compartment association (‘PC1’, the first component of a PCA on the Hi-C correlation matrix) accumulated progressively during reprogramming, are widespread and highly reproducible (Pearson R>0.97) (**Fig.2b, Supplementary Fig.2b**). Overall proportions assigned to A and B compartments remained unchanged throughout reprogramming, with ∼40% corresponding to A and ∼60% to B (**Supplementary Fig.2c**). Compartmentalization strength (as measured by average contact enrichment within and between compartments), however, was dynamically altered. OSKM induction initially (D2-D4) strengthened A-B compartment segregation, followed by substantial compartmentalization loss due to reduced contact frequencies within the B compartment and increased inter-compartment contacts (D6 onwards) (**Supplementary Fig.2d**).

**Figure 2.**
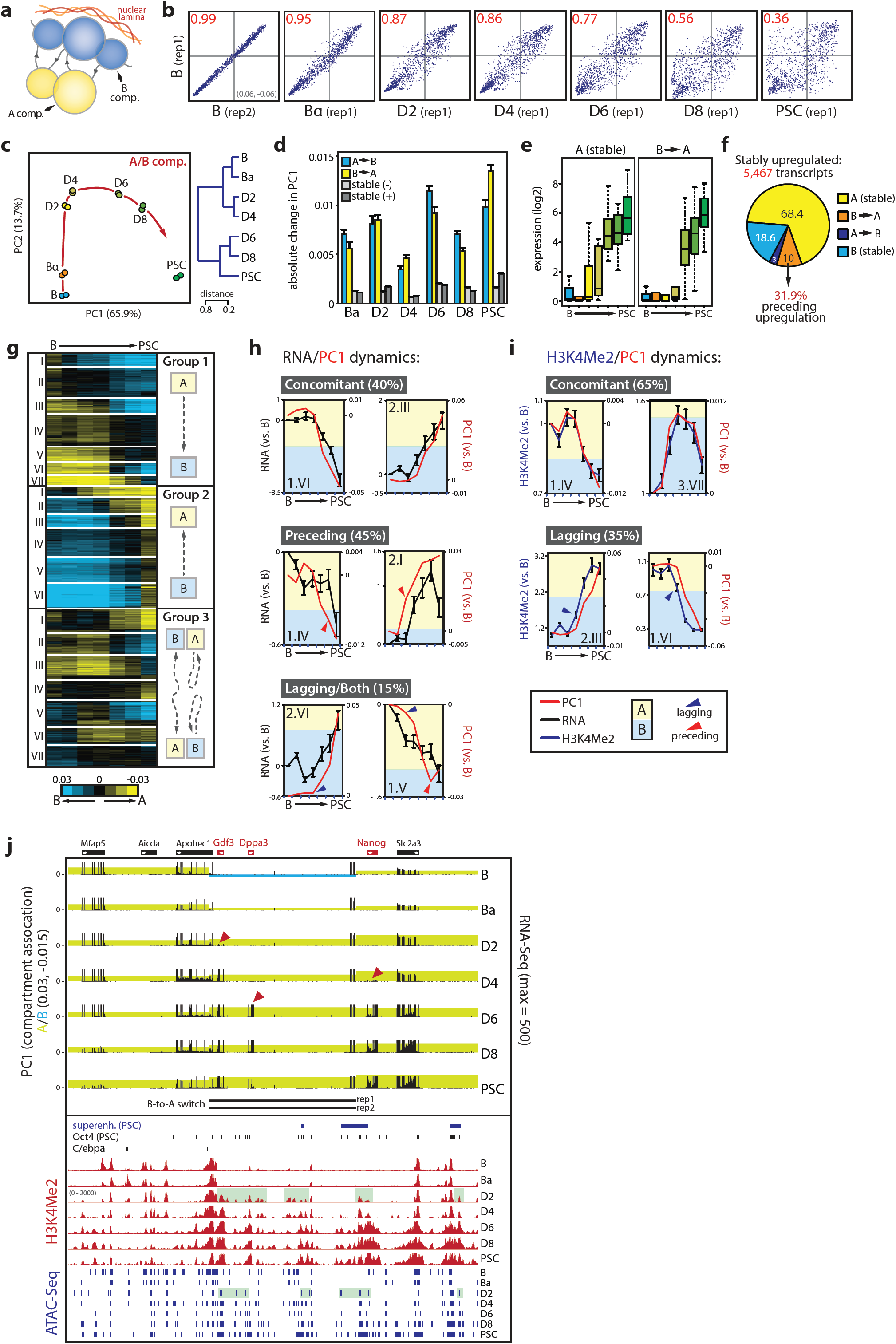
Kinetics of subnuclear compartmentalization, the transcriptome and epigenome. (**a**) Schematic representation of chromosome compartments. (**b**) Scatterplots of PC1 values (100kb bins) showing quantitative changes to the initial B cell compartmentalization during reprogramming for chromosome 13. Pearson correlation coefficient (R^2^) is indicated in red. (**c**) Principal component analysis (left, red arrow indicates hypothetical trajectory) and unsupervised hierarchical clustering (right) of PC1 values that define A/B compartments. (**d**) Absolute PC1 changes per timepoint for genomic regions that switch compartment or do not switch (‘stable’) but increase (-) or decrease (+) in PC1 value. (**e**) Box plots of gene expression kinetics (normalized counts) for key pluripotency genes (n=25) that are stably associated with the A compartment (left) or switch from A to B (right). (**f**) Compartment switching at stably upregulated genes during reprogramming. (**g**) K-means clustering (k=20) of PC1 values for 100kb genomic bins that switch compartment at any timepoint. (**h**) Examples of individual switching clusters with concomitant gene expression and PC1 changes (top, 8/20), clusters with PC1 changes preceding expression changes (middle, 9/20), and clusters (bottom) with expression changes preceding PC1 changes (1/20) or with both phenomena (2/20). (**i**) Examples of individual switching clusters that show concomitant PC1/K4Me2 changes (top, 13/20) or K4Me2 kinetics preceding PC1 modulation (bottom, 7/20). (**j**) Genome browser view of the Gdf3-Dppa3-Nanog locus. Top part shows integrated PC1 (yellow/blue shading denoting A/B compartment status, scale is indicated on the side) and RNA-Seq values (black, scale is indicated on the side), with B-to-A switch regions per replicate indicated by black bars below. Red arrowheads denote activation timing of relevant pluripotency genes. Bottom part depicts superenhancer (SE) location, Oct4 binding, C/EBPα binding as well as H3K4Me2 dynamics (red) and ATAC-Seq peaks (dark blue) during the time course. Note priming of Dppa3/Nanog enhancers at D2 (green shading). Error bars in the figure represent SEM.

Switching of loci between the A/B compartments was frequent, with 20% of the genome changing compartment at any time point during reprogramming. B-to-A and A-to-B switching each occurred in 10% of the genome, with 35% of these regions being involved in multiple switching events (**Supplementary Fig.2e**). PCA analysis revealed a reprogramming trajectory of genome compartmentalization highly similar to that seen for the transcriptome (**Fig.2c, Supplementary Fig.2f**), suggesting that both processes are dynamically coupled throughout cell fate conversion. Genes that stably switch compartment after reprogramming tend to change expression accordingly (**Supplementary Fig.2g**), confirming previous observations^38^. A lineage-specific signature existed among these genes: A-to-B switching genes were associated with immune system processes (e.g. B cell specification factor Ebf1), while B-to-A switching genes were enriched for early developmental functions (e.g. naïve pluripotency gene Tfcp2l1) (**Supplementary Fig.2h**). Compartment switching typically occurred in regions with low PC1 values at the edges of A or B domains, including regions enriched for sub-megabase compartment domains (**Supplementary Fig.2i-j**). At any time point, regions that switched also displayed the most substantial PC1 changes, suggesting that loci with a less pronounced compartment association are more amenable to changing their compartment status (**Fig.2d, Supplementary Fig.2k**).

Our dataset allows us to monitor genome architecture and to study its interplay with chromatin state and gene expression changes over time. The core transcriptional network that defines B cell identity^39^ (**Supplementary Table 2**) resided primarily (88%) in the A compartment (e.g. Ebf1, Pax5, Foxo1), of which 32% switched to B (**Supplementary Fig.3a**). Both switching and non-switching genes were rapidly silenced, but switching genes were repressed to a larger extent. In contrast, 40% of core pluripotency genes^40^ (**Supplementary Table 2**) initially resided in the B compartment of which 90% switched to A (**Supplementary Fig.3b**). Pluripotency genes already in the A compartment were activated early (D2-D4, e.g., Oct4), while genes that underwent switching from B-to-A were activated late (D6, e.g. Sox2) (**Fig.2e**). We next divided all genes that change expression between endpoints (>0.5 log2 absolute fold change) into stable (non-switching) and compartment-switching groups. As seen for the core cell identity programs, downregulated genes that changed compartment from A-to-B (21%) were silenced to a greater extent than non-switching genes in A (**Supplementary Fig.3c**). Likewise, upregulated genes that switched from B-to-A (16%) were upregulated more substantially than genes already residing in A, albeit with slower kinetics. Moreover, quantitative changes in compartment association occurred before transcriptional upregulation (**Supplementary Fig.3d**). To further explore whether compartment switching can precede transcriptional changes we examined four clusters of genes (5,467 in total) stably upregulated at early, intermediate or late time points (**Supplementary Fig.3e**). Nearly a third of the genes (175/548) that switch from B-to-A in these clusters did so before being upregulated (**Fig.2f, Supplementary Fig.3f**). Moreover, genes associated with PSC-SEs showed a significant increase in A compartment association at D2 prior to transcriptional upregulation at D4 (**Supplementary Fig.3g, Fig.1h**).

**Figure 3.**
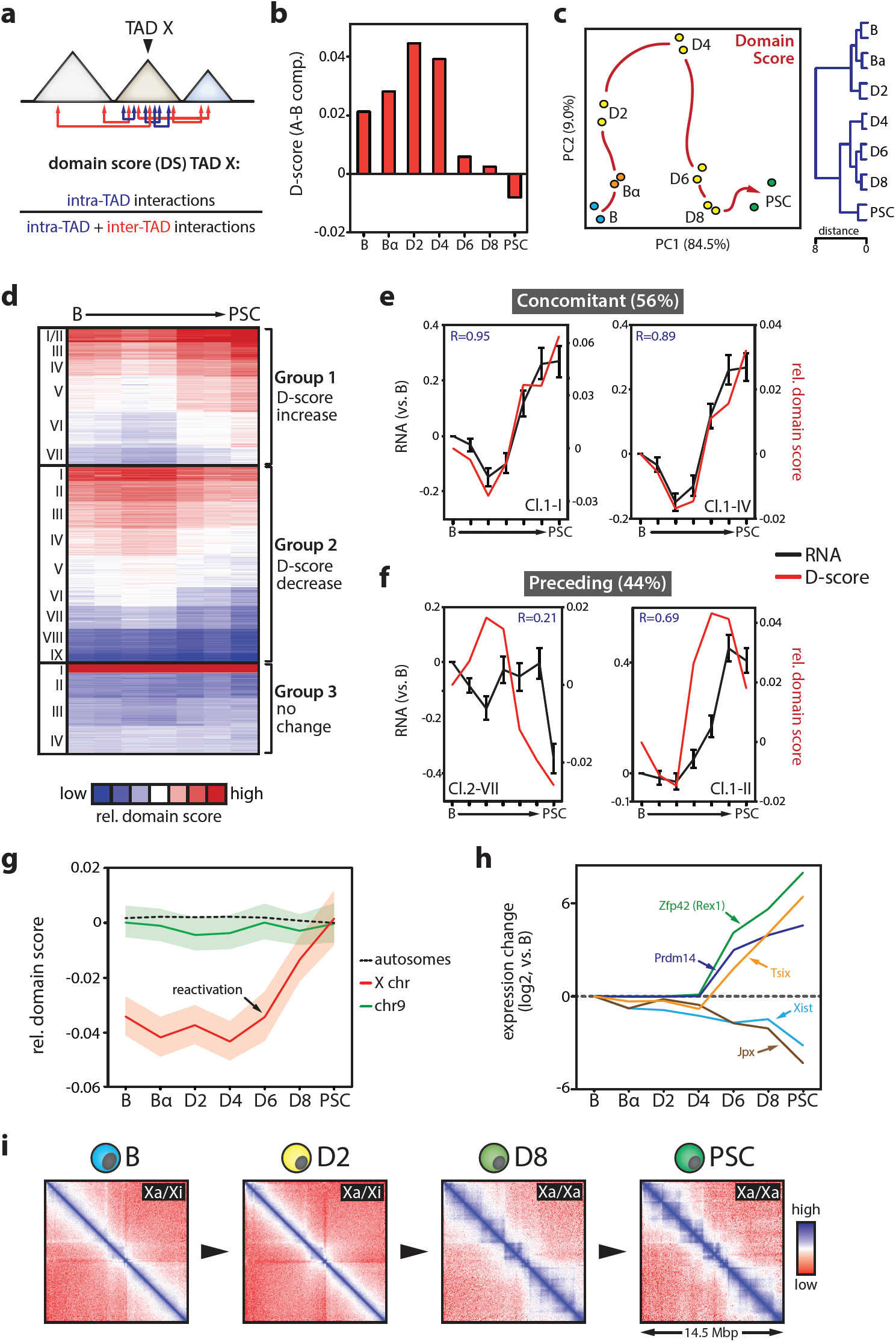
Dynamics of topologically associated domains (TADs) during reprogramming. (**a**) Cartoon depicting domain score (D-score) calculation. (**b**) Difference in D-score between TADs in the A and B compartment for each timepoint during reprogramming. (**c**) Principal component analysis (left, red arrow indicates hypothetical trajectory) and unsupervised hierarchical clustering (right) of D-score kinetics. (**d**) K-means clustering (k=20) of genome-wide relative D-score (centered on mean). (**e**) Examples of individual dynamic (16/20) D-score clusters for which gene expression and D-Score kinetics are synchronous (representing 56% of the clusters) or (**f**) where DS changes precede transcriptional changes (44%, error bars show SEM). R-values denote Pearson correlation coefficients. (**g**) Average relative D-score changes for chromosome 9, all autosomes combined and the X chromosome (left, shading denotes 95% CI). (**h**) Gene expression changes (versus B cells) of key regulators of X-chromosome re/inactivation during reprogramming. (**i**) In-situ Hi-C contact maps (50kb resolution) of a 14.5 Mb region on the X chromosome in B-D2 cells carrying one inactive X (Xi)/one active X (Xa) and D8-PSC cells carrying two Xa.

We performed K-means clustering on the 20% of the genome (n=8218 genes) that switches compartment during reprogramming, revealing 20 clusters with a wide range of switching dynamics that included non-linear and abortive trajectories (**Fig.2g**). Eight of the 20 clusters displayed concomitant changes in compartmentalization and gene expression (R>0.9, **Fig.2h**). The remainder, although generally also showing strong correlations between gene expression and PC1 (average R=0.86, range: 0.56-0.97) (**Supplementary Fig.3h**), consisted of clusters with at least one time point at which this correlation was lost (**Fig.2h**). Genes in these clusters were enriched for metabolic and secretory functions, as well as developmental processes (**Supplementary Fig.3i**). Strikingly, 9 of the 20 clusters showed changes in subnuclear compartment status preceding changes in transcriptional output (e.g. cluster 2.I, **Fig.2h**). In only a single cluster compartment topological modification lagged behind changes in gene expression, while 2 of the 20 clusters displayed both preceding and lagging relationships (**Fig.2h, Supplementary Fig.3h**). We furthermore observed a very strong overall correlation between chromatin state dynamics (gain or loss of H3K4Me2) and genome compartmentalization (average R=0.95, range: 0.93-0.98), with concomitant changes in H3K4Me2 levels and gene expression occurring in 13 of the 20 clusters. However, in 7 of the 20 clusters H3K4Me2 dynamics preceded PC1 changes (**Fig.2i**), implicating chromatin state as a driver of subnuclear compartmentalization. The extended Nanog locus provides a prime example of modifications to compartmentalization and chromatin state preceding transcriptional changes. It includes a region encompassing Gdf3, Dppa3 and the -45kb Nanog SE^37,41^, which already switched from the B to the A compartment in Bα cells. OSKM induction strengthened A compartment association of the entire locus, activated Gdf3 expression and primed the Nanog and Dppa3 regulatory elements (H3K4Me2/ATAC+) at D2 for subsequent gene activation at D4 and D6 (**Fig.2j**).

These data show that genome compartmentalization and chromatin state are dynamically reorganized during cell fate conversion and that both are tightly coupled to global changes in gene expression. In addition, a sizable number of genes are subject to changes in compartmentalization before expression alterations.

### A plastic genome architecture is acquired late in reprogramming

We used chromosome-wide insulation potential to identify TAD borders and define TADs^42^, detecting ∼3000 highly reproducible borders per time point (**Supplementary Fig.4a-b**). We analyzed differences in general TAD features at the different time points by determining the overall impact of TAD structure on gene expression (see Supplemental Materials). TADs explained a greater proportion of expression variability than linear neighborhoods. However, this proportion was progressively reduced during reprogramming (**Supplementary Fig.4c**). To measure the connectivity of a given TAD, we computed a domain score (D-score) defined by the ratio of intra-TAD interactions over all *cis* interactions (**Fig.3a**)^38^. D-scores positively correlated with gene expression and A compartment association (**Supplementary Fig.4d-e**), as previously noted^38,43^. At D4, when cells started to acquire a pluripotent phenotype, average D-scores per TAD increased (**Supplementary Fig.4f**). The positive correlation between D-scores, gene expression and compartment association seen at early time points was progressively weakened after D4 (**Fig.3b, Supplementary Fig.4d-e**), suggesting that genome architecture becomes less topologically rigid at late stages of reprogramming. Together with the observed reduced overall A-B compartment segregation (**Supplementary Fig.2d**) these data suggest that at the topological level cells gradually acquire a plastic state characteristic of the pluripotent genome^44^.

**Figure 4.**
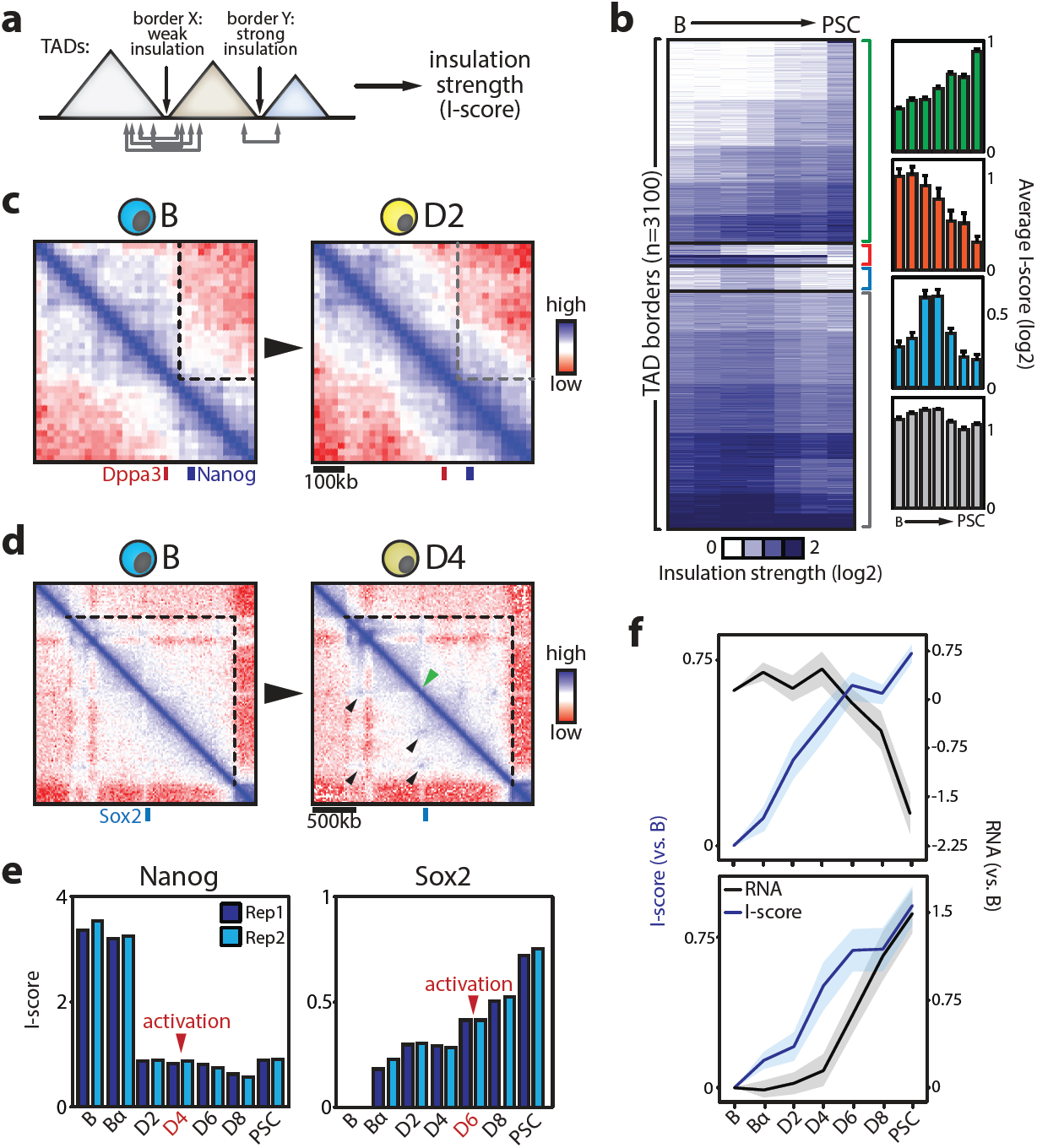
Kinetics of domain insulation during reprogramming. (**a**) Cartoon illustrating the concept of the insulation strength score (I-score). (**b**) K-means clustering (k=20) of I-score. Bar graphs show average values per timepoint for the four different categories. (**c**) In-situ Hi-C contact maps (20kb resolution) of the Dppa3-Nanog border comparing B and D2 timepoints or (**d**) the internal Sox2 border comparing B and D4 timepoints (black arrows indicate loop formation, green arrow indicates new border). (**e**) I-score kinetics of the Nanog and Sox2 borders for both replicate experiments. (**f**) Gene expression kinetics at the most dynamic border regions (n=184) where I-score changes precede transcriptional modulation (49%, n=85). Error bars and line graph shading in the figure represent 95% CI.

### Altered TAD connectivity can precede transcriptional changes

PCA analysis of D-score kinetics revealed a reprogramming trajectory for TADs similar to those for compartmentalization, transcription and active chromatin (**Fig.3c**). K-means clustering showed that 79% of TADs exhibited D-score changes (i.e. >20% change compared to B cells) (**Fig.3d**). D-score kinetics correlated closely with compartmentalization (PC1) changes (R>0.84, **Supplementary Fig.4i**). Also, the most dynamic TADs frequently switched compartment and harbored genes enriched for immune cell and stem cell related functions (**Supplementary Fig.4g-h**). TADs were highly biased in their compartment association: 88% of TADs that showed a rapid increase in D-scores initially localized to the B compartment, while 83% of the TADs with substantial D-score reductions initially resided in the A compartment (**Supplementary Fig.4g**).

To assess the correlation between TAD connectivity and gene expression, we compared D-score with intra-TAD gene expression kinetics for the 16 dynamic D-score clusters (**Fig.3d**). In 9 of 16 clusters D-score changes coincided with gene expression alterations (**Fig.3e**), in particular for TADs that showed both increased D-scores and intra-TAD expression (R=0.78). However, 7 of 16 clusters showed D-score changes preceding transcriptional changes, with no clusters showing the opposite pattern (**Fig.3f**). Thus, changes in TAD connectivity frequently precede intra-TAD transcriptional modulation.

### X chromosome reactivation evokes TAD reorganization

X chromosome reactivation in PSCs is a classic model for studying the relationship between chromosome structure and gene expression^45^. The B cells used were derived from female mice carrying one inactive X chromosome, allowing us to study this process using our dataset. While average TAD connectivity for each time point remained similar on autosomes, X chromosome TADs displayed substantial gains in D-scores after D4 (**Fig.3g**). The observed chromosome-wide D-score increase is due to a reactivation of the largely TAD-devoid inactive X chromosome^11,46-48^. Indeed, after D4 TAD structures were fully re-established and key regulators of X reactivation activated (Zfp42, Prdm14, Tsix), while X chromosome repressors (Xist and Jpx) were down-regulated (**Fig.3h-i**).

### Changes in TAD border strength occur early in reprogramming

Partitioning of the genome into TADs was largely stable during reprogramming as 75%-87% of TAD borders were invariant (**Supplementary Fig.5b**). Nevertheless, we also observed the formation or loss of borders during reprogramming, resulting in a net increase in the number of borders and a reduction of average TAD size from 891kb to 741kb (**Supplementary Fig.5a-c**). TAD borders were enriched for CTCF binding sites and transcription start sites (**Supplementary Fig.5d**), as expected^17,49^. To quantify TAD border dynamics we aligned borders throughout the time course and calculated an insulation strength score (I-score) as a measure of a given border’s ability to insulate adjacent TADs^42,50^ (**Fig.4a**). PCA analysis of I-score kinetics revealed a reprogramming trajectory resembling the transcriptome, PC1, H3K4Me2 and D-score trajectories determined before (**Supplementary Fig.5e-f**). CTCF occupancy correlated with I-score, although even borders lacking CTCF (<12%) showed progressively higher I-scores during reprogramming (**Supplementary Fig.5g**). Half of all borders showed a >20% difference in I-score during reprogramming (**Fig.4b**). Meta-border plots projecting multiple borders into a single average plot confirmed that I-score dynamics reflect actual contact maps (**Supplementary Fig.6a**).

**Figure 5.**
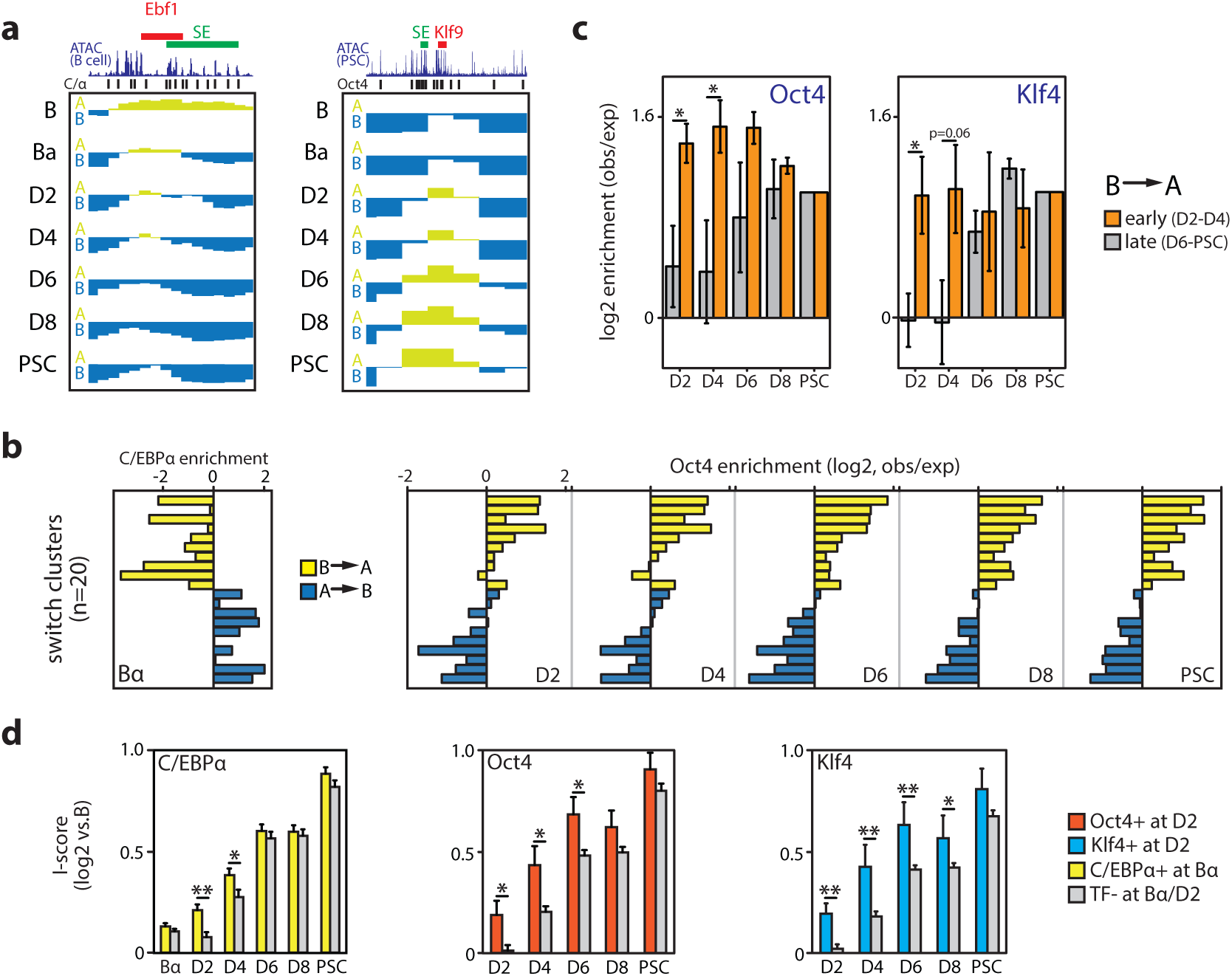
Transcription factor binding and topological genome dynamics. (**a**) Examples of C/EBPα-mediated A-to-B switching (Ebf1 locus) and OSKM-mediated B-to-A switching (Klf9). Genes are shown as red bars, superenhancers (SE) as green bars. (**b**) C/EBPα (ChIP-Seq) and Oct4 (inferred from ATAC-Seq, see Supplemental Materials) binding enrichment (over the genome-wide average) at the 20 switching clusters shown in **Fig.2g**. (**c**) Oct4 and Klf4 binding enrichment in clusters that switch B-to-A compartment early (D2-D4) or late (D6-PSC). Error bars denote SEM (*p<0.05, unpaired two-tailed *t*-test). (**d**) Insulation strength (I-score) dynamics at hyper-dynamic borders (n=184) bound or not bound by the indicated transcription factors (TFs). Error bars denote SEM (*p<0.05, **p<0.01; unpaired two-tailed *t*-test).

We determined how I-score dynamics correlated with gene regulatory processes and found a highly significant representation of genes with cell type-specific functions (e.g. immune system, developmental biology) at border regions, in addition to the expected housekeeping genes^17^ (**Supplementary Fig.6b**). Pluripotency genes were often found at or near border regions (**Supplementary Fig.6c**), including Nanog and Sox2. Both of these loci showed rapid I-score changes that preceded their transcriptional activation (**Fig.4c-e**). In B and Bα cells Nanog was separated from Dppa3 by a strong border in a region that harbors the -45kb Nanog SE and Gdf3 (**Fig.4c, Fig.2j**), probably preventing the spatial clustering of these genes and enhancers in PSCs^51^. The I-score was considerably reduced at D2 after OSKM induction (**Fig.4e**), facilitating interactions between genes and their enhancers required for subsequent transcriptional activation (D4-D6). Indeed, 4C-Seq analyses showed increased cross-border contact frequencies of the Nanog promoter as early as D2 (**Supplementary Fig.6d**). Within the Sox2-TAD, a new internal border and several chromatin loops appeared between Bα and D4 stages that progressively isolated Sox2 together with its essential downstream SE^52,53^, an event likely necessary for Sox2 activation at D6 (**Fig.4d-e, Supplementary Fig.6e**).

To further understand the relationship between I-score changes and the expression of nearby genes we analyzed transcriptional changes at the 184 most dynamic borders regions that increase in insulation strength (>75% change in I-score). Gene expression was altered at many of these borders (46%) during reprogramming, with no clear bias for activation or repression. At 49% of these borders (n=43/88) I-scores increased before transcriptional changes (**Fig.4f**), while for the remaining borders a mix of concomitant (n=15), lagging (n=15) and more complex (n=15) kinetics was observed. Likewise, I-score changes also preceded modulation of chromatin state and subnuclear compartmentalization (**Supplementary Fig.6f-g**). Thus, altered insulation strength at TAD borders is an early reprogramming event linked to transcriptome re-wiring.

### Transcription factors drive topological genome reorganization

Since lineage-instructive TFs have the capacity to modify genome architecture^24,54^, we investigated the impact of C/EBPα and OSKM on genome topology. Approximately 5% of the genome switched compartment during the C/EBPα-induced B-to-Bα transition and 5% during the OSKM-induced Bα-to-D2 transition. Of these early switching regions, only 29% (B-to-Bα) and 36% (Bα-to-D2) represented stable switches (**Supplementary Fig.7a**). C/EBPα had a largely repressive effect (66% A-to-B switching, e.g. Ebf1), while OSKM operated predominately as an activator (70% B-to-A switches, e.g. Klf9) (**Fig.5a, Supplementary Fig.7a**). Both C/EBPα and OSKM evoked A-to-B switching and transcriptional silencing of B cell-related loci. At D2, OSKM induced B-to-A switching and activation of known target genes of pluripotency factors involved in developmental processes (**Supplementary Fig.7b**). However, genes undergoing stable B-to-A switching in Bα cells were only upregulated after OSKM activation, including genes implicated in early embryonic development such as Gdf3 and Dppa3 (**Supplementary Fig.7c**). Globally, C/EBPα binding was strongly enriched in the previously identified A-to-B switching clusters and depleted in B-to-A switching clusters (**Fig.5b**). In contrast, Oct4 and Klf4 binding (as inferred by ATAC-Seq) was concentrated in B-to-A switching regions (**Fig.5b, Supplementary Fig.7d**). This biased genomic distribution was already apparent at D2 and was stably maintained or reinforced, with early switching clusters (D2-D4) being rapidly targeted by Oct4 and Klf4 and late switching clusters (D6-PSC) becoming more gradually enriched (**Fig.5c**).

We next examined TF action at TAD borders. Oct4 target sites within ∼30% of all border regions were already accessible at D2 (**Supplementary Fig.7e**). Oct4 or Klf4 recruitment to the most dynamic borders at D2 correlated with accelerated I-score gains as compared to borders bound at later time points (**Fig.5d, Supplementary Fig.7f**). C/EBPα-bound borders increased their I-scores more rapidly only after OSKM activation at D2 (**Fig.5d**) and Oct4 enrichment was significantly higher at borders previously bound by C/EBPα (**Supplementary Fig.7g**), suggesting that C/EBPα primes border regions for subsequent OSKM-induced topological changes. Moreover, Oct4, Klf4 and C/EBPα were frequently recruited to the same dynamic borders early in reprogramming (**Supplementary Fig.7h**).

TF-bound sites cluster over large distances^14,54-56^. We therefore addressed the dynamics of such 3D crosstalk during reprogramming by measuring inter-TAD spatial connectivity between TF-bound genomic sites (within a 2-10 Mb window, analogous to PE-SCAn^54^). Leveraging our dense contact maps (at 5kb resolution), we observed strong interactions between Ebf1 or Pu.1 binding sites in B cells in agreement with their function as key B cell regulators (**Fig.6a**). These interaction networks largely disappeared for Ebf1 in Bα and for Pu.1 in D4 cells. Spatial clustering of C/EBPα targets was already present in B cells (**Fig.6a**), indicating that C/EBPα exploits existing 3D interaction hubs, such as those formed by Pu.1. Alongside interaction hubs mediated by hematopoietic TFs, Oct4 binding sites clustered from D2 onwards to establish 3D crosstalk between PSC-specific regulatory elements (**Fig.6a**), showing that topological interaction hubs mediated by lineage-specific and pluripotency TFs can coexist. Moreover, Nanog targeted regions formed interaction hubs as early as D2, before the gene becomes expressed at D4 (**Fig.6a**), suggesting that late pluripotency factors hitchhike onto an OSKM-mediated interaction hub to lock-in the PSC fate.

**Figure 6.**
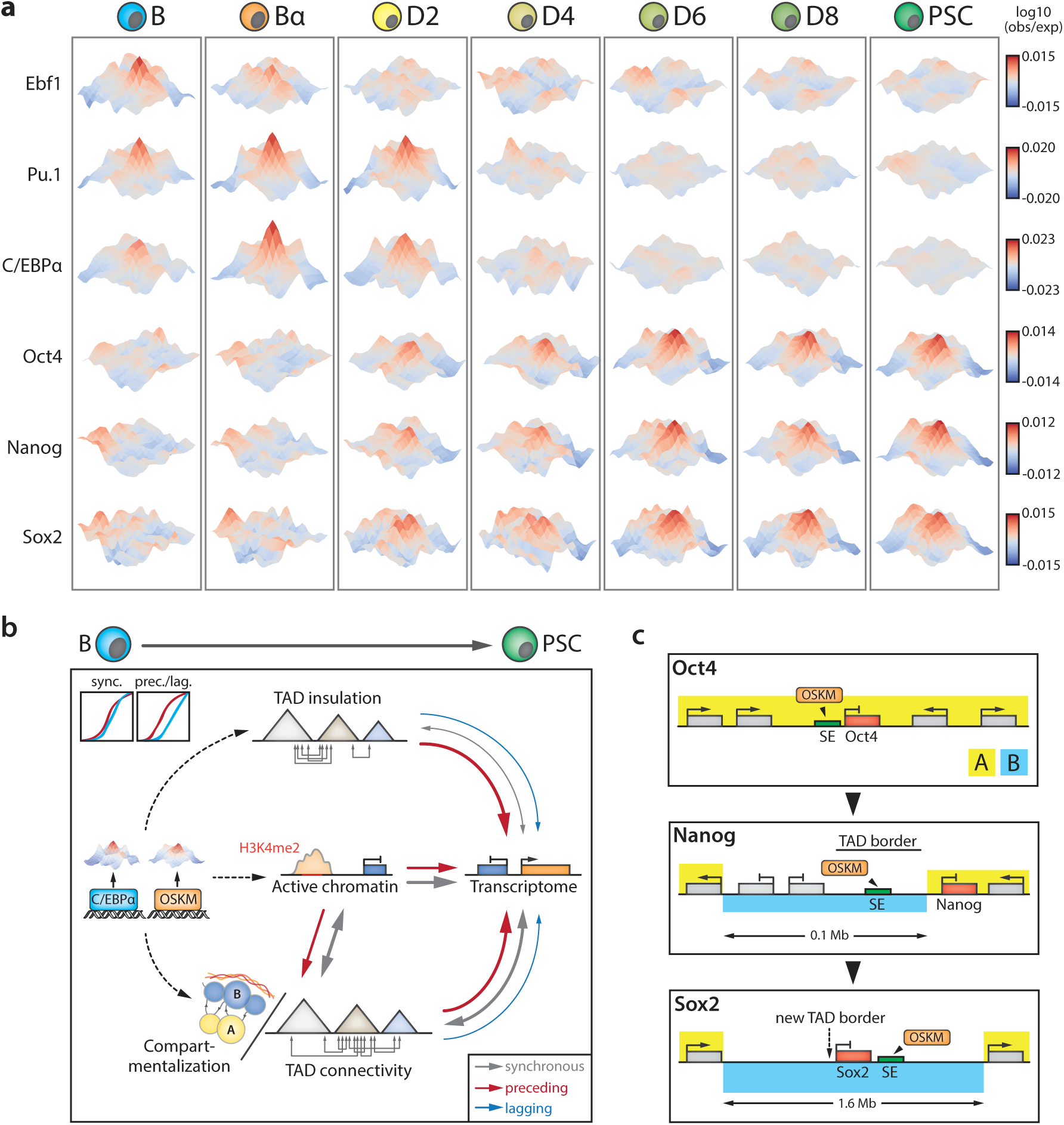
Dynamics of 3D crosstalk between transcription factor target sites and model schemes. (**a**) 3D interaction meta-plots (5kb resolution) depicting interaction frequencies of sites bound by the indicated TFs during reprogramming. Hubs visualize inter-TAD crosstalk between TF binding sites 2-10 Mb apart. Area shown is centered on the respective TF binding sites (+/- 50kb). (**b**) Schematic depicting the interplay between lineage-instructive transcription factors (TFs), chromatin state, genome topology and gene regulation during cell reprogramming. Arrows denote functional relationships of a synchronous (gray), preceding (red) or lagging (blue) nature (illustrated by the line graph inset), with arrow thickness indicating prevalence. Successive introduction of C/EBPα and OSKM in B cells induces (dashed arrows), possibly via the formation of long-range interaction hubs, a topological reorganization of compartments and TADs that enables gene expression changes for conversion to pluripotent stem cells (PSC). TFs can also operate indirectly via changes in chromatin state, the latter appearing to be a major driver of compartmentalization and TAD connectivity dynamics. Note that transcriptional changes preceding modifications to topology also occur, but at much lower frequencies. (**c**) Activation scenarios for the pluripotency factors Oct4, Nanog and Sox2. Oct4 activation does not seem to require major topological modifications, as the gene and its superenhancer (SE) already reside in the A compartment in B cells and TAD border strength is unaltered. Nanog activation is preceded by B-to-A compartment switching of its nearby SE as well as a decrease in TAD border strength that facilitates Nanog-SE interaction. Sox2 activation is preceded by the formation of a new TAD border through chromatin loop formation that insulates the gene and its SE; concomitant with activation of the gene the complete 1.6 Mb region switches from the B to the A compartment.

In summary, C/EBPα and OSKM binding correlates with accelerated topological remodeling of compartmentalization and TAD insulation. In addition, computing inter-TAD 3D crosstalk between TF targets enabled us to visualize the stage-specific formation and disassembly of these interaction hubs during reprogramming.

## DISCUSSION

Our analysis of somatic cell reprogramming (summarized in **Supplementary Fig.7i**) revealed that the overall dynamics of genome topology, chromatin state and gene expression are closely coupled. Nevertheless, much of this coupling occurs in a non-synchronous manner: changes in subnuclear compartmentalization, TAD connectivity and TAD insulation frequently precede transcriptional changes, with the reverse situation occurring only at low frequencies. We propose that TFs induce successive changes in genome architecture to enable gene regulatory rewiring during cell reprogramming (**Fig.6b**). The observed stepwise increase in the accessibility of Oct4 binding sites further implies that TFs encounter new binding opportunities as topological and regulatory landscapes evolve. Overall, our data indicates that genome topology represents an instructive force harnessed by TFs for implementing transcriptional changes and that this is a general principle of cell fate decisions.

Our findings also provide an explanation for the sequential activation of the pluripotency factors Oct4, Nanog and Sox2 in spite of the cells’ continuous exposure to the Yamanaka factors (**Fig.6c**). The embedding of Oct4 and its enhancers within an A compartment domain, surrounded by genes highly expressed in B cells, may explain its almost immediate activation by OSKM without detectable topological alterations. In contrast, the late activation of Nanog and Sox2 is preceded and accompanied by substantial changes in compartmentalization and TAD structure, indicating that the removal of topological barriers creates new opportunities for gene regulation. That active chromatin dynamics often anticipate changes in subnuclear compartmentalization suggests it plays a major role in mediating switches between the active A and the inactive B compartments (**Fig.6b**), in line with imaging and local chromatin conformation analyses^57,58^. The progressive loss of architectural rigidity seen during reprogramming is consistent with reduced organization of inactive chromatin in PSCs^54^. Experiments aimed at perturbing specific topological changes and testing their effect on cell fate represent a next frontier in dissecting the relationship between genome form and function.

How do TFs drive 3D genome changes? C/EBPα and Oct4 are selectively enriched in different regions destined to switch compartment. Here, TFs could act by inducing the subnuclear repositioning of specific loci^59^, for example by initiating modification of local chromatin states. In addition, the TFs rapidly induce insulation strength changes at the most dynamic TAD borders, independent of major changes in compartmentalization or chromatin state. Separate modes of action for TFs at these two topological levels seem plausible, as compartmentalization and TAD organization have been suggested to depend on distinct mechanisms^60,61^. The inter-TAD hubs of TF target regions likely contribute to compartment reorganization and suggest that TFs can exploit topologies previously established by other TFs. As early targets, SE regions may provide key platforms for TFs to achieve topological genome remodeling^62^. The ability of lineage instructive regulators to alter genome topology raises the possibility that they possess unappreciated architectural functions at distinct topological layers.

## ACKNOWLEDGEMENTS

We thank D. Higgs, J. Hughes, J. Davies and Z. Duan for advice on Hi-C technology; C. Schmidl for ChIPm-Seq advice; C. van Oevelen for help with CTCF ChIP-Seq.; C. Segura for mouse colony management and T. Tian for bone marrow collection; the CRG Genomics Core Facility and the CRG-CNAG Sequencing Unit for sequencing and Graf laboratory members for discussions. This work was supported by the European Research Council Synergy Grant (4D-Genome) and Ministerio de Educacion y Ciencia, SAF.2012-37167. R.S. was supported by an EMBO Long-term Fellowship (ALTF 1201-2014) and a Marie Curie Individual Fellowship (H2020-MSCA-IF-2014).

## AUTHOR CONTRIBUTIONS

R.S. and T.G. conceived the study and wrote the manuscript with input from all co-authors. R.S. performed molecular biology, RNA-Seq, ChIP(m)-Seq, ATAC-Seq, 4C-Seq and in-situ Hi-C experiments. R.S, E.V., F.S, J.Q., A.G., S.C. and M.A.M-R. performed bioinformatic analyses. R.S., E.V., F.S. and M.A.M-R. integrated and visualized data. B.D.S. performed reprogramming experiments with help from R.S. and C.B. R.S., F.L.D. and Y.C. optimized and implemented in-situ Hi-C technology. J.H. performed high-throughput sequencing. F.L.D., G.F., M.B. and M.A.M-R. provided valuable advice and T.G. supervised the research.

## METHODS

### Mice

We crossed ‘reprogrammable mice’ containing a doxycycline-inducible OSKM cassette and the tetracycline transactivator^63^ with an Oct4-GFP reporter strain^64^, as previously described^29,30^. B cells were isolated from 8 to 16 week old female mice. Mice were housed in standard cages under 12h light–dark cycles and fed ad libitum with a standard chow diet. All experiments were approved by the Ethics Committee of the Barcelona Biomedical Research Park (PRBB) and performed according to Spanish and European legislation.

### Cell culture & somatic cell reprogramming

Embryonic stem cells (E14TG2a) and short-term induced PSCs were cultured on gelatinized plates or Mitomycin C inactivated mouse embryonic fibroblasts (MEFs) in N2B27 medium (50% DMEM-F12, 50% Neurobasal medium, N2 (100x), B27 (50x)) supplemented with small-molecule inhibitors PD (1μM, PD0325901) and CHIR (3 μM, CHIR99021), as well as LIF (10 ng ml^-1^). Reprogramming of primary B cells isolated from the bone marrow of reprogrammable/Oct4-GFP mice was performed as previously described^30^. Two independent biological replicate reprogramming experiments were used for data generation. Briefly, pre-B cells were infected with C/EBPαER-hCD4 retrovirus, plated at 500 cells cm^-2^ in gelatinized 12 well plates on Mitomycin C inactivated MEF feeders in RPMI medium. C/EBPα was activated by adding 100 nM β-estradiol (E2) for 18 hours. After E2 washout, the cultures were switched to N2B27 medium supplemented with IL-4 (10 ng ml^-1^), IL-7 (10 ng ml^-1^) and IL-15 (2 ng ml^-1^). OSKM was activated by adding 2 μg ml^-1^ of doxycycline. Harvesting was done at indicated time points by trypsinization followed by a 20 min pre-plating step to remove feeder cells. All cell lines have been routinely tested for mycoplasma contamination.

### RNA isolation, quantitative RT-PCR and RNA-Sequencing (RNA-Seq)

RNA was extracted using the miRNeasy mini kit (Qiagen) and quantified by Nanodrop. cDNA was produced with the High Capacity RNA-to-cDNA kit (Applied Biosystems) and used for qRT-PCR analysis in triplicate reactions with the SYBR Green QPCR Master Mix (Applied Biosystems). Primers are available upon request. Libraries were prepared using the Illumina TruSeq Stranded mRNA Library Preparation Kit followed by paired-end sequencing (2x125bp) on an Illumina HiSeq2500.

### Assay for Transposase-Accessible Chromatin with high throughput sequencing (ATAC-Seq)

ATAC-seq was performed as previously described^30^. 100,000 cells were washed once with 100 μl PBS and resuspended in 50 μl lysis buffer (10 mM Tris-HCl pH 7.4, 10 mM NaCl, 3 mM MgCl2, 0.2% IGEPAL CA-630). Cells were centrifuged for 10 min at 500g (4°C), supernatant was removed and nuclei were resuspended in 50 μl transposition reaction mix (25 μl TD buffer, 2.5 μl Tn5 transposase and 22.5 μl nuclease-free water) and incubated at 37°C for 45 min. DNA was isolated using the MinElute DNA Purification Kit (Qiagen). Library amplification was performed by two sequential PCR reactions (8 and 5 cycles, respectively). Library quality was checked on a Bioanalyzer, followed by paired-end sequencing (2x75bp) on an Illumina HiSeq2500.

### Chromatin Immunoprecipitation followed by high throughput sequencing (ChIP[m]-Seq)

ChIP-Seq using tagmentation (ChIPm-Seq) was performed as previously described^34^ with 100,000 crosslinked cells using 1 μl of H3K4me2 antibody (Abcam, ab32356) per IP. Tagmentation of immobilized H3K4me2-enriched chromatin was performed for 2 min at 37°C in 25 μl transposition reaction mix (12.5 μl TD buffer, 1.0 μl Tn5 transposase and 11.5 μl nuclease-free water). Library amplification was performed as described for ATAC-Seq. Library quality was checked on a Bioanalyzer, followed by sequencing (1x75bp) on an Illumina NextSeq500. Conventional ChIP-Seq was performed as previously described^65^ with 300,000 crosslinked cells using 5 μl of CTCF antibody (Millipore, 07-729). Libraries were prepared using the Illumina TruSeq ChIP Library Preparation Kit and sequenced (1x50bp) on an Illumina HiSeq2500.

### Chromosome Conformation Capture followed by high-throughput sequencing (4C-Seq)

4C-seq was performed as described previously^66,67^. Briefly, 0.5-1.0 million crosslinked nuclei were digested with Csp6I followed by ligation under dilute conditions. After decrosslinking and DNA purification, samples were digested overnight with DpnII and once more ligated under dilute conditions. Column-purified DNA was directly used as input for inverse PCR using primers (available upon request) with Illumina adapter sequences as overhangs. Several PCR reactions were pooled, purified and sequenced (1x75bp) on an Illumina HiSeq2500.

### Gene Ontology (GO) analysis

GO analyses were performed using the Molecular Signatures Database (MSigDB)^68^ for gene lists or GREAT^69^ for peak lists. Only statistically significant (FDR<0.01) terms and pathways were used.

### In-situ Hi-C library preparation

In-situ Hi-C was performed as described^11^ with the following modifications: 1) Two million cells were used as starting material; 2) chromatin was initially digested with 100 U MboI (New England Biolabs) for 2 hours, followed by addition of another 100U (2 hour incubation) and a final 100U before overnight incubation; 3) before fill-in with bio-dATP, nuclei were pelleted and resuspended in fresh 1x NEB2 buffer; 4) ligation was performed overnight at 24°C using 10,000 cohesive end units per reaction; 5) decrosslinked and purified DNA was sonicated to an average size of 300-400 bp using a Bioruptor Pico (Diagenode; 7 cycles of 20 s on and 60 s off); 6) DNA fragment size selection was only performed after final library amplification; 7) library preparation was performed with the NEBNext DNA Library Prep Kit (New England Biolabs) using 3 μl NEBNext adaptor in the ligation step; 8) libraries were amplified for 8-12 cycles using Herculase II Fusion DNA Polymerase (Agilent) and purified/size-selected using Agencourt AMPure XP beads (>200 bp). Hi-C Library quality was assessed by ClaI digest and low-coverage sequencing on an Illumina NextSeq500, after which every technical replicate (n=2) of each biological replicate (n=2) was sequenced at high-coverage on an Illumina HiSeq2500. Data from technical replicates was pooled for downstream analysis. We sequenced >18 billion reads in total to obtain 0.78-1.21 billion valid interactions per timepoint per biological replicate (see **Supplementary Table 1** for dataset statistics).

### Gene expression analysis using RNA-Seq data

Reads were mapped using STAR^70^ (-outFilterMultimapNmax 1 -outFilterMismatchNmax 999 - outFilterMismatchNoverLmax 0.06 -sjdbOverhang 100 –outFilterType BySJout - alignSJoverhangMin 8 -alignSJDBoverhangMin 1 -alignIntronMin 20 -alignIntronMax 1000000 - alignMatesGapMax 1000000) and the Ensembl mouse genome annotation (GRCm38.78). Gene expression was quantified using STAR (--quantMode GeneCounts). Sample scaling and statistical analysis were performed using the R package DESeq2^71^ (R 3.1.0 and Bioconductor 3.0) and vsd counts were used for further analysis unless stated otherwise. Standard RPKM values were used as an absolute measure of gene expression. Genes changing significantly at any time point were identified using the nbinomLRT test (FDR<0.01) and for>2-fold change between at least two time points (average of two biological replicates, vsd values). Clustering was performed using the Rpackage Mfuzz (2.26.0).

### Chromatin accessibility analysis using ATAC-Seq data

Reads were mapped to the UCSC mouse genome build (mm10) using Bowtie2^72^ with standard settings. Reads mapping to multiple locations in the genome were removed using SAMtools^73^; PCR duplicates were filtered using Picard (http://broadinstitute.github.io/picard). Bam files were parsed to HOMER^74^ for downstream analyses and browser visualization. Peaks in ATAC-Seq signal were identified using *findPeaks* (-region -localSize 50000 -size 250 -minDist 500 - fragLength 0, FDR<0.001).

### ChIP(m)-Seq data analysis

Reads were mapped and filtered as described for ATAC-Seq. H3K4me2 enriched regions were identified using HOMER *findpeaks* (findPeaks -region -size 1000 -minDist 2500, using a mock IgG experiment as background signal). H3K4me2 coverage per 100kb genomic bin was computed using BEDTools^75^ and normalized for differences in sequencing depth (normalized coverage = coverage / (number of unique mapped reads in dataset / 1e6)). CTCF peaks were identified using MACS2^76^ (https://github.com/taoliu/MACS/) with *callpeak* --nolambda --nomodel -g mm -- extsize 100 -q 0.01.

### 4C-Seq data analysis

The sequence of the 4C-Seq reading primer was trimmed from the 5’ of reads using the demultiplex.py script from the R package fourCseq^77^ (allowing 4 mismatches). Reads in which this sequence could not be found were discarded. Reads were mapped using STAR and processed using fourCseq to filter out reads not located at the end of a valid fragment and to count reads per fragment. Signal tracks were made after smoothing RPKM counts per fragment with a running mean over five fragments.

### In-situ Hi-C data processing and normalization

We processed Hi-C data using an in-house pipeline based on TADbit^78^. First, quality of the reads was checked using FastQC (http://www.bioinformatics.babraham.ac.uk/projects/fastqc/) to discard problematic samples and detect systematic artifacts. Trimmomatic^79^ with the recommended parameters for paired end reads was used to remove adapter sequences and poor quality reads (ILLUMINACLIP:TruSeq3-PE.fa:2:30:12:1:true; LEADING:3; TRAILING:3; MAXINFO:targetLength:0.999; and MINLEN:36).

For mapping, a fragment-based strategy as implemented in TADbit was used, which is similar to previously published protocols^80^. Briefly, each side of the sequenced read was mapped in full length to the reference genome (mm10, Dec 2011 GRCm38). After this step, if a read was not uniquely mapped, we assumed the read was chimeric due to ligation of several DNA fragments. We next searched for ligation sites, discarding those reads in which no ligation site was found. Remaining reads were split as often as ligation sites were found. Individual split read fragments were then mapped independently. These steps were repeated for each read in the input FASTQ files. Multiple fragments from a single uniquely mapped read will result in as many contact as possible pairs can be made between the fragments. For example, if a single read was mapped through three fragments, a total of three contacts (all-versus-all) was represented in the final contact matrix. We used the TADbit filtering module to remove non-informative contacts and to create contact matrices. The different categories of filtered reads applied are:

- *self-circle*: reads coming from a single restriction enzyme (RE) fragment and point to the outside.
- *dangling-end*: reads coming from a single RE fragment and point to the inside.
- *error*: reads coming from a single RE fragment and point in the same direction.
- *extra dangling-end*: reads coming from different RE fragments but are close enough and point to the inside. The distance threshold used was left to 500 bp (default), which is between percentile 95 and 99 of average fragment lengths.
- *duplicated*: the combination of the start positions and directions of the reads was repeated, pointing at a PCR artifact. This filter only removed extra copies of the original pair.
- *random breaks*: start position of one of the reads was too far from RE cutting site, possibly due to non-canonical enzymatic activity or random physical breaks. Threshold was set to 750 bp (default), > percentile 99.9.

From the resulting contact matrices, low quality bins (those presenting low contacts numbers) were removed as implemented in TADbit’s “filter_columns” routine. A single round of ICE normalization ^81^ - also known as “vanilla” normalization ^11^ - was performed. That is, each cell in the Hi-C matrix was divided by the product of the interactions in its columns and the interactions in its row. Finally, all matrices were corrected to achieve an average content of one interaction per cell.

### Identification of subnuclear compartments and topologically associated domains (TADs)

To segment the genome into A/B compartments, normalized Hi-C matrices at 100kb resolution were corrected for decay as previously published, grouping diagonals when signal-to-noise was below 0.05^11^. Corrected matrices were the split into chromosomal matrices and transformed into correlation matrices using the Pearson product-moment correlation. The first component of a PCA (PC1) on each of these matrices was used as a quantitative measure of compartmentalization and H3K4Me2 ChIPm-Seq data was used to assign negative and positive PC1 categories to the correct compartments. If necessary, the sign of the PC1 (which is randomly assigned) was inverted so that positive PC1 values corresponded to A compartment regions and vice versa for the B compartment.

Normalized contact matrices at 50kb resolution were used to define TADs, using a previously described method with default parameters^42,46^. First, for each bin, an insulation index was obtained based on the number of contacts between bins on each side of a given bin. Differences in insulation index between both sides of the bin were computed and borders were called searching for minima within the insulation index. The insulation score of each border was determined as previously described^42^, using the difference in the delta vector between the local maximum to the left and local minimum to the right of the boundary bin. This procedure resulted in a set of borders for each time point and replicate. To obtain a set of consensus borders along the time course, we proceeded in two steps: (a) merging borders of replicates and overlapping merged borders (that is, for each pair of replicates, we expand borders one bin on each side and kept only those borders present in both replicates as merged borders), and (b) we further expanded two extra bins (100kb) on each side and determined the overlap to get a consensus set of borders common to any pair of time points.

Domain scores were obtained by averaging cells over parts of the Hi-C matrix. In nature, this metric is sensitive to outlier cells with a lot of counts and is less sensitive to missing data. For this analysis (and for the meta-loop analysis below) we thus used a more stringent strategy to remove low-coverage bins by fitting a logistic function to the distribution of the sum of interactions in each bin:

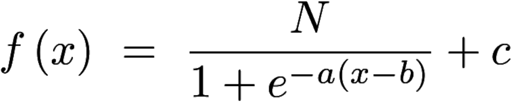

Where f is the logistic function optimized by the variables a, b and c. N is the number of bins in the matrix, and x the number of interactions in a given bin. This fit was implemented by weighting bins with higher values of interactions, as we considered bins with lower counts artifacts. We set the weight function as dependent on the bin index, in the context of bins sorted by their sum of interactions:

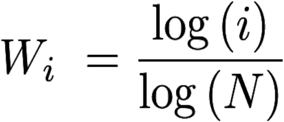

With i representing the index of the bin and W the weight applied to the fitting. Once the logistic function was fitted, we used it to define a threshold. We removed bins with fewer counts than x when f(x) was equal to zero. The resulting filtered matrices were ICE normalized (1 round, see above). Finally, domain scores were calculated using matrices binned at 50kb by dividing the sum of intra-TAD contacts by the sum of all contacts involving the TAD.

### Expression variability explained by TADs

To estimate expression variability, we fitted a hierarchical regression model per gene expression values for each timepoint, including three levels of organization: the gene itself, the local neighborhood (the 50 kb TSS bin) and the TAD. We used the variance associated with each level and the total variance of the model to assess the proportion of variability explained by each factor. In order to test if topology was playing a role beyond linear proximity of genes, we repeated the estimation replacing actual TADs by a fixed segmentation of the genome in domains with the same size as the average TAD (i.e. “fake” TADs, constructed by placing a border at fixed distances that correspond to the average size of TADs). Model estimation was performed using the lme4 R package.

### Inter and intra-compartment strength measurements

We followed a previously reported strategy to measure overall interaction strengths within and between A and B compartments^60^. Briefly, we based our analysis on the 100kb bins showing the most extreme PC1 values, discretizing them by percentiles and taking the bottom 20% as B compartment and the top 20% as A compartment. We classified each bin in the genome according to PC1 percentiles and gathered contacts between each category, computing the log2 enrichment over the expected counts by distance decay. Finally, we summarize each type of interaction (A-A, B-B and A-B/B-A) by taking the median values of the log2 contact enrichment.

### Meta-analysis of borders and interactions between transcription factor binding sites

To assess whether particular parts of the Hi-C interaction matrices had common structural features, we performed meta-analyses by merging individual sub-matrices into an average meta-matrix in a similar fashion as previously published^54^. Two types of meta-analysis were performed. First, we studied TAD border dynamics at 50kb resolution by extracting interaction counts 1.25Mb up and downstream of the TAD border. Extracted matrices were averaged for each group of clustered TAD borders, including those that increase, decrease or do not change in insulation score during reprogramming. Second, we studied whether two regions bound by a given transcription factor (TF) are likely to find each other more than expected within a genomic distance ranging from 2 to 10Mb. All sub-matrices at 5kb resolution between pairs of TF binding sites and 50kb up and downstream of a TF peak were extracted and averaged into a single meta-matrix. For Oct4, Nanog and Sox2 meta-analyses we used those TF binding sites that overlapped with an ATAC-Seq peak (see above) at the D2 stage. All meta-analyses were performed using the observed/expected Hi-C matrices, which were filtered, ICE normalized and corrected for decay. For visualization proposes, the resulting meta-analysis matrices were smoothed using a Gaussian filter of sigma=1.

### Data availability

All data generated has been deposited in the Gene Expression Omnibus (GEO) under GSE96611. Accession number of published datasets used: CTCF ChIP-Seq in pre-B cells: SRR397837^82^; CTCF ChIP-Seq in induced PSCs: GSE76478^38^; Oct4 and Nanog ChIP-Seq in PSCs: GSE44286^37^; Klf4 ChIP-Seq in PSCs: GSE11431^83^; C/EBPα and Pu.1 ChIP-Seq in Bα cells: GSE71215^30^; Ebf1 V5-ChIP-Seq in pro-B cells: GSE53595^84^.

**Supplementary Figure 1.**
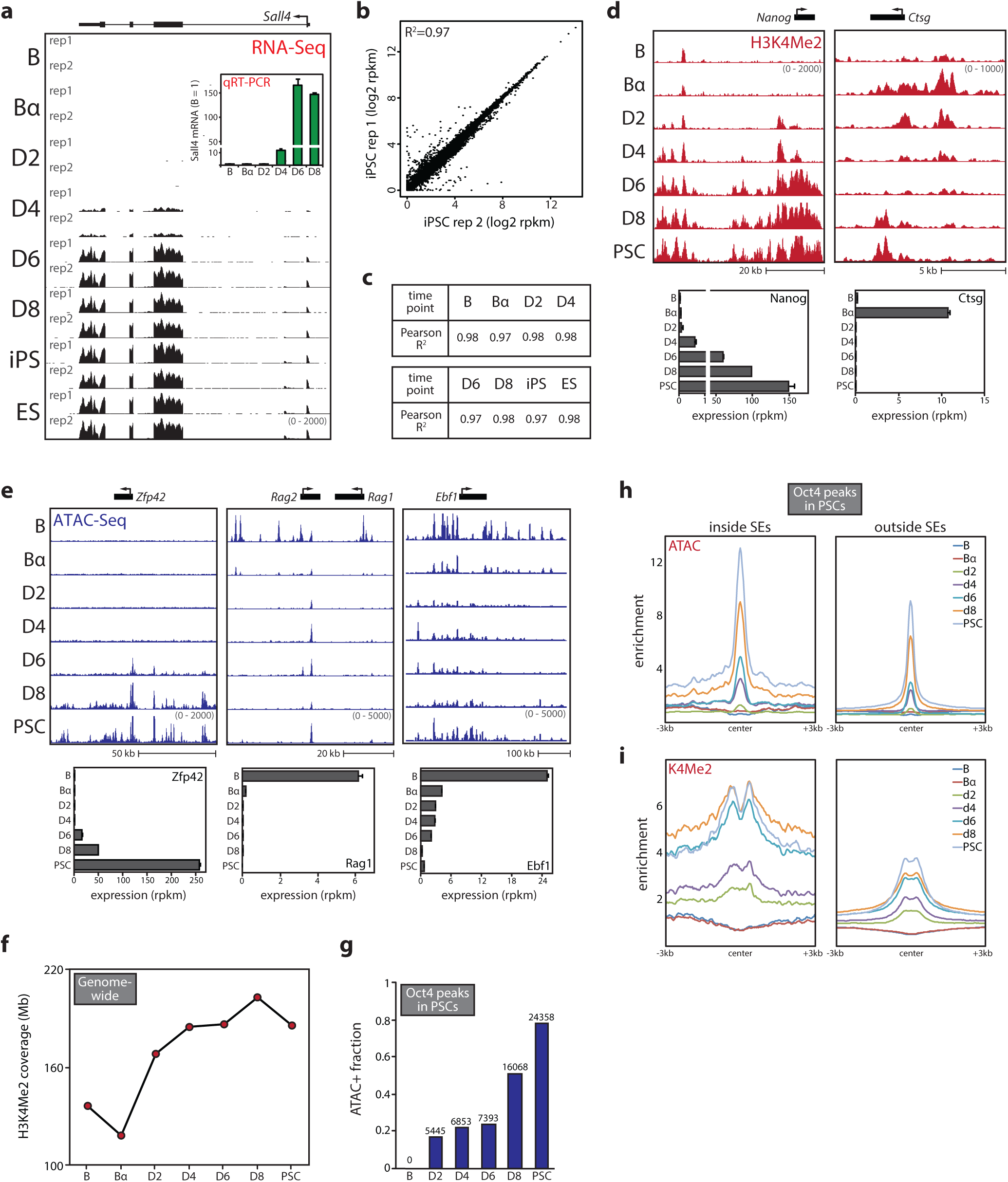
Transcriptome and epigenome dynamics during reprogramming. (**a**)Genome browser view of Sall4 gene expression measured by RNA-Seq data (two biological replicates per timepoint). Bar graph insert depicts qRT-PCR measurements of Sall4 expression. (**b**)Scatterplot of RPKM gene expression values for biological replicates 1 and 2 (iPS samples shown). (**c**) Pearson correlation (R^2^) values between RNA-Seq replicates for all timepoints. (**d**) Genome browser views of the Nanog and Ctsg genes with H3K4Me2 ChIPm-Seq profiles during reprogramming. Bar graphs show gene expression dynamics for these genes. (**e**) Genome browser views of the Zfp42, Rag1-Rag2 and Ebf1 genes with ATAC-Seq profiles during reprogramming. Bar graphs show gene expression dynamics for these genes. (**f**) Normalized genome-wide H3K4Me2 (marking active chromatin) coverage per timepoint. (**g**) Fraction of Oct4 binding sites in PSCs overlapping with an ATAC-Seq peak (‘ATAC+’) during reprogramming. Absolute numbers of sites are shown above each bar. (**h**) ATAC-Seq and (**i**) H3K4Me2 coverage profiles for Oct4 binding sites in PSCs inside (left) and outside (right) superenhancers (SEs) during reprogramming. Error bars in the figure denote 95% CI.

**Supplementary Figure 2.**
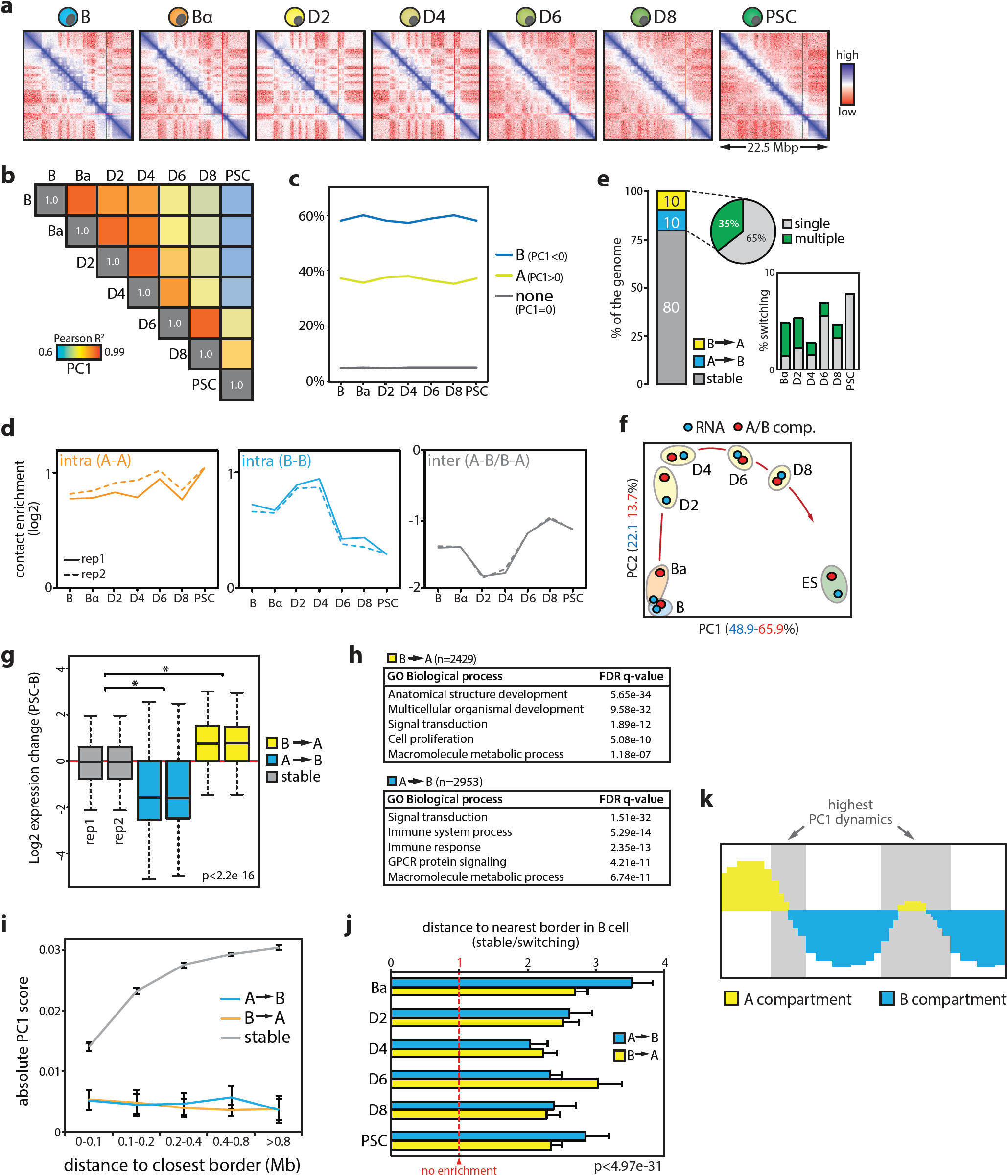
Subnuclear compartmentalization dynamics during reprogramming. (**a**) Example in-situ Hi-C contact maps (50kb resolution) of a 22.5 Mb region on chromosome 3. (**b**) Pearson correlation coefficient (R^2^) heatmap of PC1 value comparisons between timepoints. (**c**) Line chart depicting genome fractions assigned to A or B compartments at the different time points. Regions that could not be assigned (PC1=0, e.g. telomeric regions) are shown in gray. (**d**) Overall contact enrichment for 100kb bins within the A (left) or B (middle) compartment or between A and B (right) compartments during reprogramming. (**e**) Fraction of the genome that switches compartment at any point during the time course. Bar graph depicts switching percentages per timepoint. (**f**) Scaled overlay of principal component analyses for gene expression (blue) and compartmentalization (red) dynamics. (**g**) Gene expression changes for genes in bins that switch compartment at any timepoint or do not switch (‘stable’) during reprogramming. *p<2.2e-16, Wilcoxon rank-sum test. (**h**) Gene ontology terms associated with the two categories of switching genes. (**i**) Absolute PC1 score of switching or non-switching (‘stable) bins as a function of their distance to the nearest compartment border. (**j**) Average distance to the nearest compartment border of non-switching stable bins divided by the average distance of the two types of switching bins. Switching bins are significantly closer to borders than stable bins at all timepoints (Poisson regression, p<4.97e-31). (**k**) Cartoon summarizing characteristics of compartment switching dynamics: compartmentalization dynamics are highest in regions of low PC1 and near compartment domain borders. Error bars in all plots denote 95% CI.

**Supplementary Figure 3.**
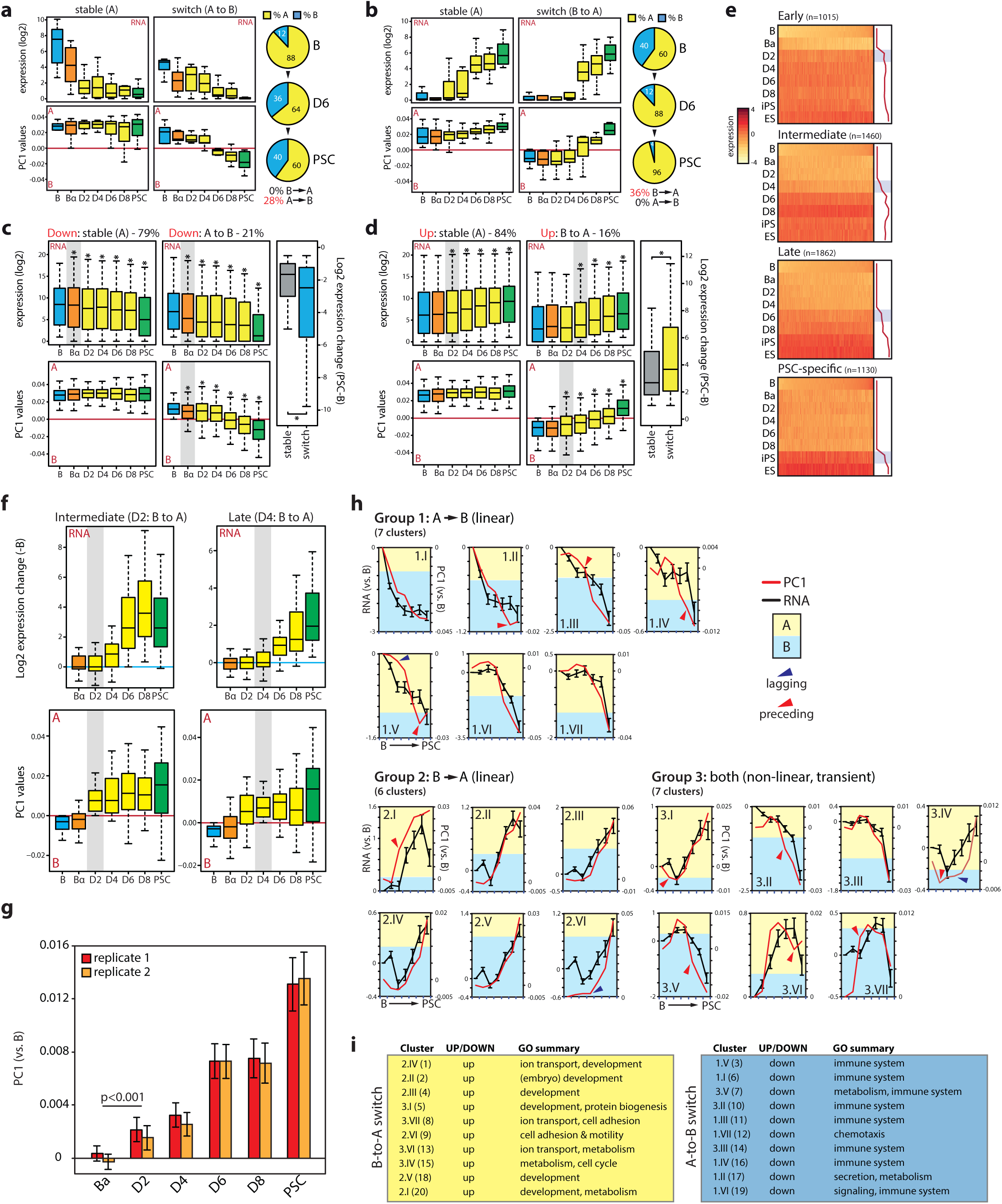
Relationship between subnuclear compartmentalization and gene expression changes. (**a,b**) Comparison of gene expression and PC1 dynamics for key B cell (panel a) and pluripotency (panel b) genes (n=25). Genes were grouped into those stably associated with the A compartment (left) and those that switch (right). Pie charts depict changes in compartment status for these genes during reprogramming. (**c,d**) Gene expression (top) and PC1 (bottom) kinetics for downregulated genes (<-0.5 log2 fold change, panel c) or upregulated genes (>0.5 log2 fold change, panel d) between reprogramming endpoints. Genes were grouped into those stably associated with the A compartment (left) and those that switch (right). Gray shading marks first timepoint of significant change (versus B, *p<0.01, Wilcoxon rank-sum test). Boxplots on the right depict the extent of expression change (PSC versus B) for the two groups of genes. (**e**) Gene expression clusters of genes stably upregulated during reprogramming at different stages. Line graphs on the right depict average kinetics, gray shading marks first timepoint of significant change. (**f**) Gene expression (top) and PC1 (bottom) kinetics for stably upregulated genes (from two clusters shown in panel e) that switch compartment preceding transcriptional upregulation. Gray shading indicates timepoint at which switching was completed. (**g**) Change in PC1 value (relative to B cells) during reprogramming for bins containing PSC superenhancers (p<0.001, unpaired two-tailed *t*-test). (**h**) Kinetics of gene expression and PC1 change for all 20 individual switching clusters (**see Fig.2g**). Arrows indicate time points were the correlation between expression and PC1 is lost. (**i**) Summarized gene ontology (GO) annotation of the 20 switching clusters. Error bars in all plots denote SEM.

**Supplementary Figure 4.**
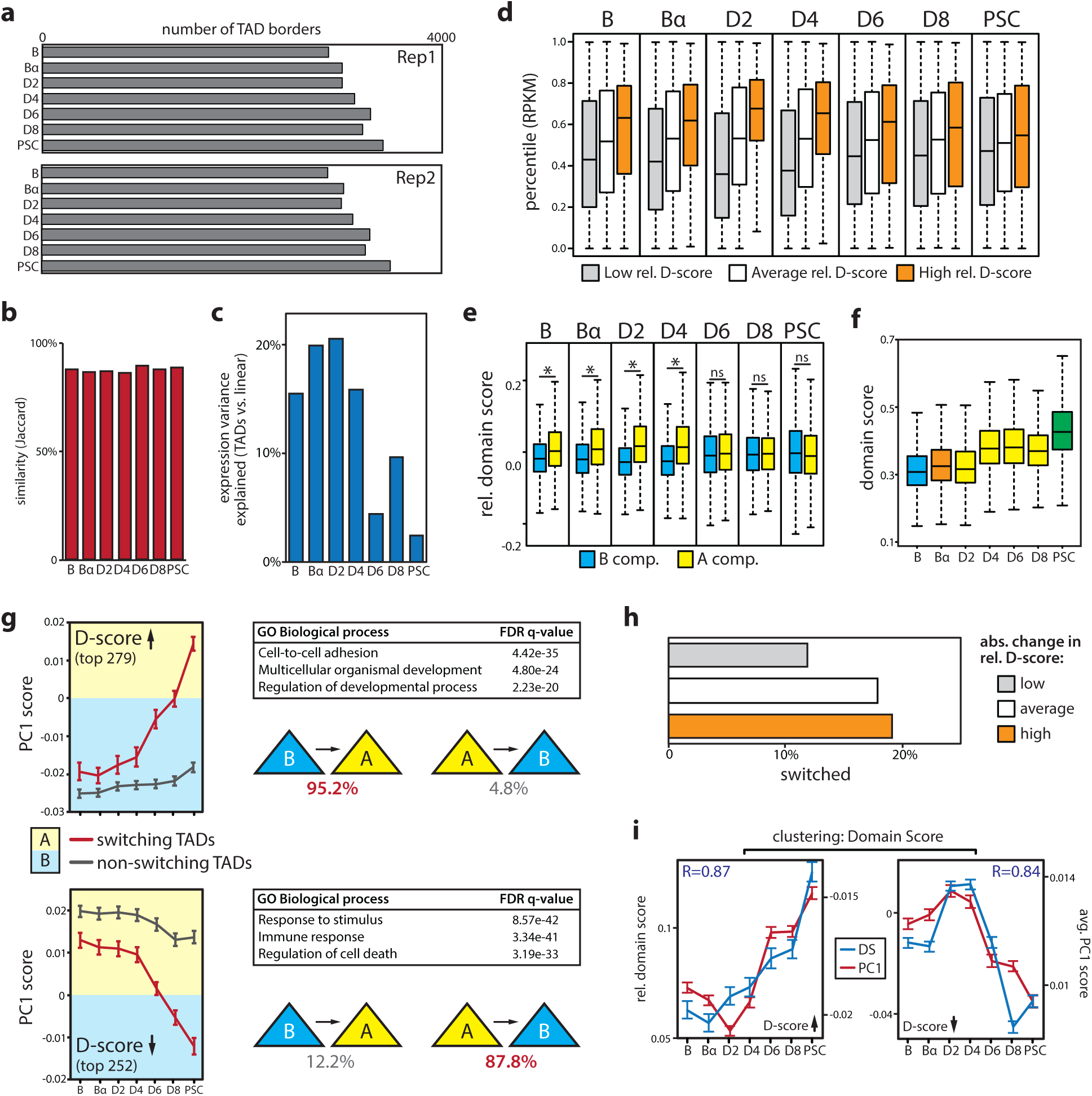
Topologically associated domain (TAD) dynamics during reprogramming. (**a**) Number of TAD borders identified per timepoint for each biological replicate. (**b**) TAD border reproducibility between replicates as measured by the Jaccard index. (**c**) Percentage of expression variance explained by TADs (relative to a linear model, see Supplemental Materials for a detailed explanation) for each timepoint. (**d**) Average expression of genes (plotted as an expression percentile) in TADs having a low (-0.26;-0.02), average (-0.02; 0.1) or high (0.1;0.6) relative domain score (D-score). (**e**) Boxplots showing relative D-score values for TADs in the A or B compartment at each timepoint. *p<0.001, Wilcoxon rank-sum test, ns = non significant. (**f**) Average genome-wide D-score during reprogramming. (**g**) Top dynamic TADs gaining (upper half) or losing (lower half) D-score during reprogramming. Line graphs show PC1 values for switching and non-switching TADs. Percentages of A-to-B and B-to-A switching for both groups of TADs are depicted by triangles. Tables show selected gene ontology (GO) terms for the genes within the corresponding TADs. (**h**) Fraction of TADs that switch compartment in groups of TADs with low (0-0.02), average (0.02-0.07) or high (>0.07) absolute changes in D-score. (**i**) Average D-score and PC1 kinetics during reprogramming for clusters of TADs that gain (left) or lose (right) D-score. Pearson correlation coefficients (R) are indicated. Error bars in all plots denote SEM.

**Supplementary Figure 5.**
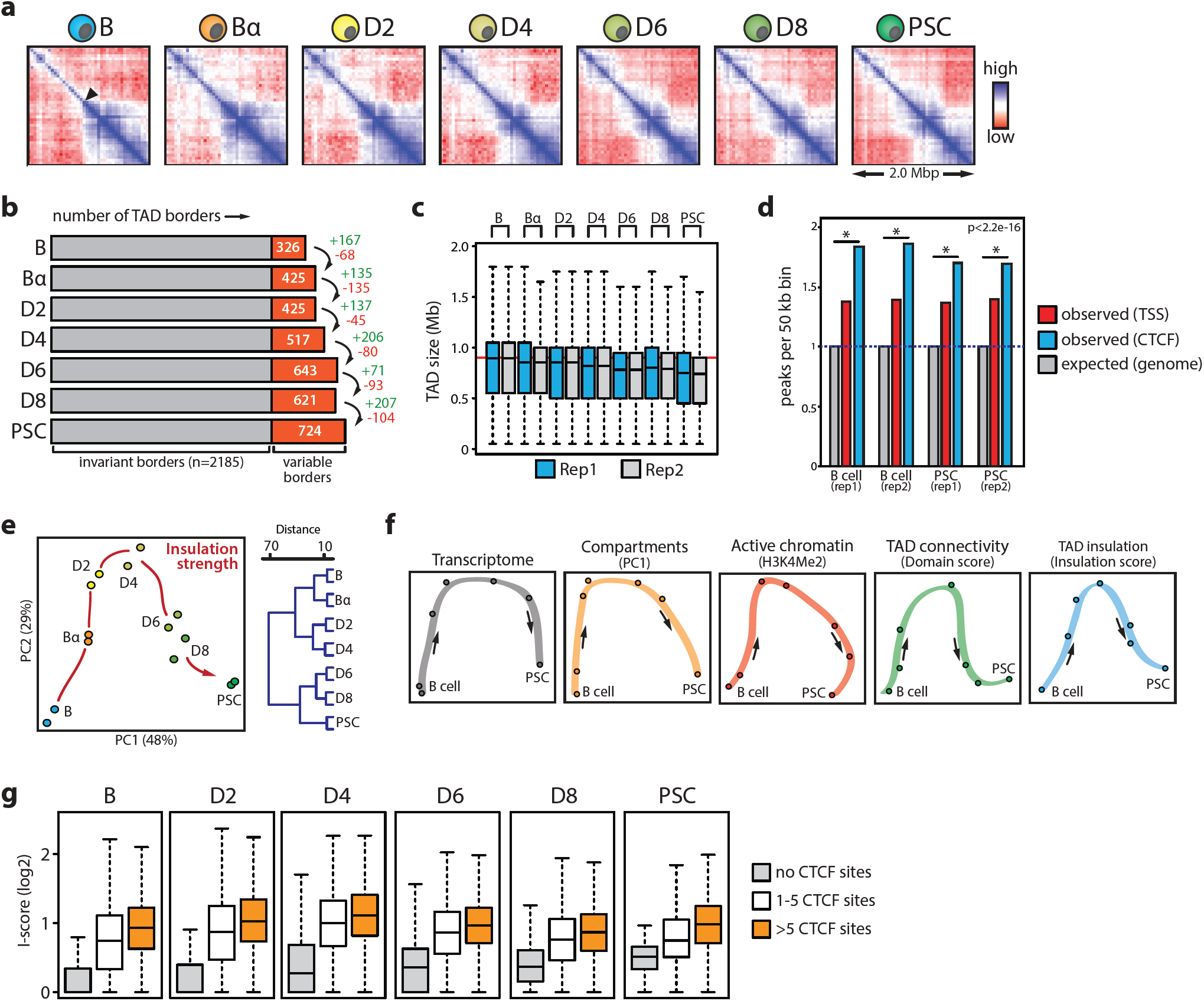
TAD border insulation strength dynamics during reprogramming. (**a**) In-situ Hi-C contact maps (50kb resolution) of a 2.0 Mb region on chromosome 14 centered on TAD border 999. (**b**) Number of TAD borders reproducibly called per timepoint. Invariant borders were present at all timepoints; variable borders were lost/acquired during reprogramming. (**c**) Boxplots showing TAD size distributions during reprogramming. (**d**) Enrichment of CTCF and transcription start sites (TTS) at borders (compared to their genome-wide distribution) in two replicate datasets for B cells and PSCs. *p<2.2e-16, Wilcoxon rank-sum test. (**e**) Principal component analysis (PCA) and unsupervised hierarchical clustering of insulation score (I-score) values. (**f**) Collection of all PCA trajectories generated in this study. Points denote average data from two biological replicates. (**g**) Boxplots depicting I-score of borders harboring no CTCF sites, 1-5 CTCF sites or >5 CTCF sites for indicated timepoints.

**Supplementary Figure 6.**
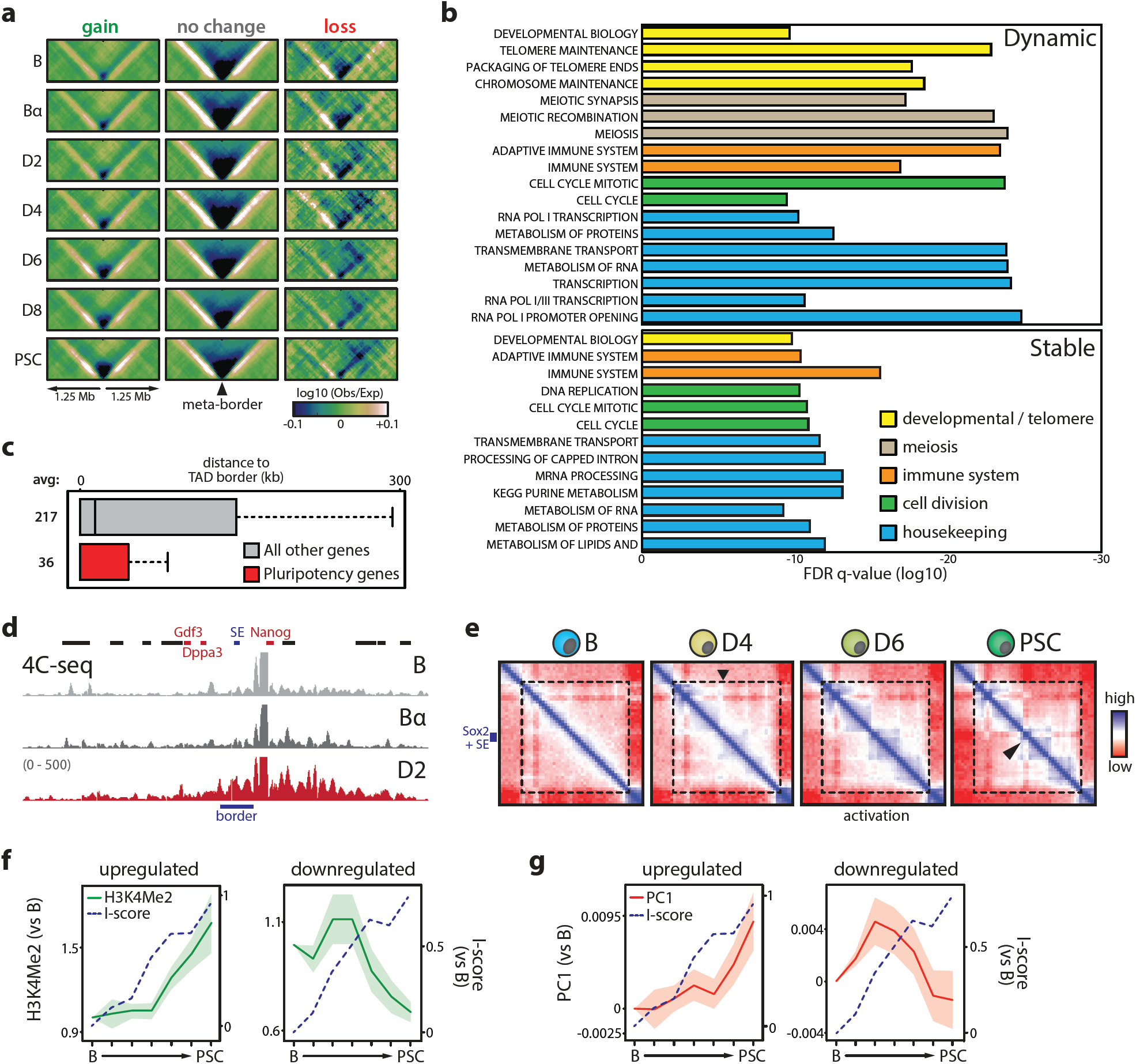
Cell type-specific genes reside near dynamic TAD borders. (**a**) Meta-border plots for all borders that gain insulation score (I-score), do not change I-score or lose I-score. (**b**) Gene ontology terms significantly associated with genes found within dynamic (top) or stable (bottom) border regions. (**c**) Boxplot showing the average distance of pluripotency genes (red) or all other genes (gray) to the nearest TAD border. (**d**) 4C-Seq analysis of the Dppa3-Nanog locus at early reprogramming timepoints using the Nanog promoter as a viewpoint (border region is indicated in blue). (**e**) 2.25 Mb in-situ Hi-C contact maps (50 kb resolution) centered on the Sox2 gene and its superenhancer (SE). The appearance of an internal border at D4 is indicated by a black arrow. Note the progressive insulation of Sox2 and its SE into a smaller domain as the gene is activated (indicated by a black arrow in the PSC map). (**f,g**) Kinetics of H3K4Me2 (panel f) and PC1 (panel g) changes at dynamic borders harboring genes that are either upregulated (left) or downregulated (right). Shading denotes SEM.

**Supplementary Figure 7.**
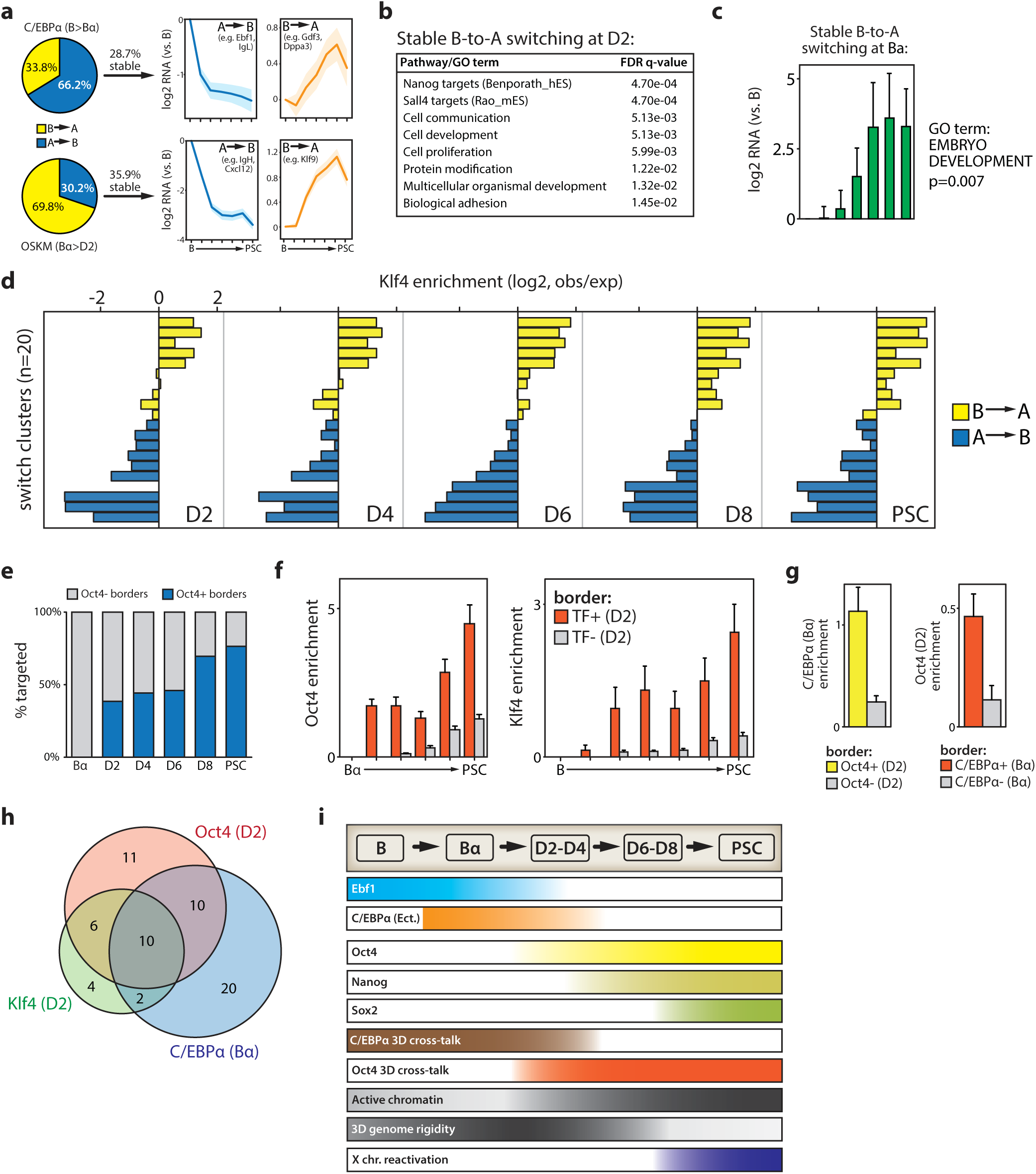
Transcription factor dynamics at hotspots of topological change. (**a**) Compartment switching induced by C/EBPα (B-to-Bα, top) or OSKM (Bα-to-D2, bottom). Line graphs depict expression changes of genes located in regions that have stably switched (shading denotes SEM). (**b**) GO annotation of genes that stably switch B-to-A compartment at the Bα-D2 transition. (**c**) Average gene expression changes of the genes associated with the gene ontology (GO) term ‘Embryo Development’ that stably switch B-to-A compartment at the B-Bα transition. (**d**) Klf4 binding enrichment (over the genome-wide average) at the 20 switching clusters shown in **Fig.2g**. (**e**) Percentage of TAD border regions bound by Oct4 at each timepoint. (**f**) Oct4 (left) and Klf4 (right) enrichment kinetics at border regions that are already targeted by these factors at D2 (red) or border regions not yet targeted at D2 (grey). (**g**) C/EBPα (left) or Oct4 (right) enrichment at border regions bound by indicated transcription factors at the earliest timepoint (yellow/red) or unbound regions (grey). (**h**) Venn diagram showing the overlap between the number of dynamic borders bound by Oct4 (at D2), Klf4 (at D2) and C/EBPα (at Bα). (**i**) Kinetics of key transcriptional, epigenomic and topological events during somatic cell reprogramming. Light-to-dark color intensity range signifies quantitative differences. Ect., ectopic; chr., chromosome

